# PSKR1 balances the plant growth-defense trade-off in the rhizosphere microbiome

**DOI:** 10.1101/2022.11.07.515115

**Authors:** Siyu Song, Zayda Morales Moreira, Xuecheng Zhang, Andrew C. Diener, Cara H. Haney

## Abstract

Microbiota provide benefits to their hosts including nutrient uptake and protection against pathogens. How hosts balance an appropriate immune response to avoid microbiota overgrowth while avoiding autoimmunity is not well understood. Here we show that *Arabidopsis pskr1* (*phytosulfokine receptor 1*) loss-of-function mutants display autoimmunity and reduced rhizosphere bacterial growth when inoculated with normally growth-promoting *Pseudomonas fluorescens*. Transcriptional profiling demonstrated that PSKR1 regulates the plant growth-defense trade-off during plant-microbiome interactions: PSKR1 upregulates plant photosynthesis and root growth but suppresses salicylic acid (SA)-mediated defense responses. Genetic epistasis experiments showed that *PSRK1* inhibition of microbiota-induced autoimmunity is fully dependent on SA signaling. Finally, using a transgenic reporter, we showed that *P. fluorescens* induces *PSKR1* expression in roots, suggesting *P. fluorescens* might manipulate plant signaling to promote its colonization. Our data demonstrate a genetic mechanism to coordinate beneficial functions of the microbiome while preventing autoimmunity.

## Introduction

Plant roots associate with diverse microbes recruited from the soil known as the rhizosphere microbiome^1,2^ that provide plants with a variety of benefits including nutrient uptake, improved plant growth, and protection from pathogens. In order to avoid overgrowth of beneficial bacteria, plants must maintain an appropriate immune response; indeed loss of plant immune signaling can result in dysbiosis and opportunistic disease from microbiota^3^. However, excessive plant defense responses can result in plant stunting and autoimmunity^4^. The fundamental question of how plants regulate the growth-defense trade-off in response to the rhizosphere microbiome is largely unexplored.

Gain of function of positive regulators or plant immunity, or loss of function of negative regulators, can result in constitutive plant immunity against foliar pathogens. Many of these hyperimmune mutants are stunted as a result of plant-growth defense tradeoffs^4^. Despite their dramatic effect on host physiology and disease resistance, described autoimmune mutants have relatively modest effects on rhizosphere microbiota^5^, or even result in increased growth of rhizosphere microbiota^6^. Whether microbiota can induce autoimmunity in plants, and the genetic mechanisms that would normally prevent this, remain to be investigated.

To identify genes that prevent autoimmunity in response to microbiota, we made use of a previous forward genetic screen that identified mutants impaired in root defense-hormone crosstalk. The *HSM* (hormone-mediated suppression of MAMP-triggered immunity) genes encode positive regulators of hormone signaling or negative regulators of pattern-triggered immunity (PTI) in roots^7^. By rescreening the *hsm* mutant collection for rhizosphere microbiome phenotypes, Song *et al*. identified 6 *hsm* mutants with higher or lower levels of rhizosphere *P. fluorescens*^8^. These include *HSM4*, which encodes the *Arabidopsis* ortholog of *SDA1*, a yeast ribosome biogenesis protein^9^, and *hsm13* that has a loss of function of *FERONIA* (*FER* kinase^8^. These findings indicate that at least some *HSM* genes may encode immune regulators required for *Arabidopsis* association with the rhizosphere microbiome.

By rescreening the *hsm* mutant collection, we identified the *hsm7* mutant that exhibits autoimmunity in response to normally growth-promoting *Pseudomonas fluorescens* inoculation. Bulk segregation and whole genome sequencing revealed that a loss-of-function of *PHYTOSULFOKINE RECEPTOR 1* (*PSKR1*) underlies the *hsm7* phenotype. The *hsm7* mutant exhibits reduced growth and increased immunity in response to microbiota indicating that PSKR1 is required for promotion of growth and inhibition of defense required to prevent autoimmunity towards the rhizosphere microbiome.

## Results

### PSKR1 prevents autoimmunity to rhizosphere microbiota

We reasoned that a plant mutant with autoimmunity against microbiota would 1) support lower levels of bacterial growth, 2) show stunted growth in response to commensal bacteria, and 3) exhibit higher levels of defense-gene expression. Previous work showed that the *hsm* mutants *hsm4, hsm6* and *hsm7* had lower rhizosphere commensal colonization levels, suggesting potential inappropriate elevated defense responses against commensal bacteria^8^. Rescreening *hsm4, hsm6* and *hsm7* by inoculating with beneficial *P. fluorescens* strain WCS365 revealed that both *hsm6* and *hsm7* had stunted root growth (Figure 1A, 1B) and that only *hsm7* showed elevated defense gene expression (Figure 1C). These results indicate that *hsm7* displays an altered plant growth-defense trade-off and autoimmunity phenotype in response to rhizosphere microbiota.

**Figure 1.**
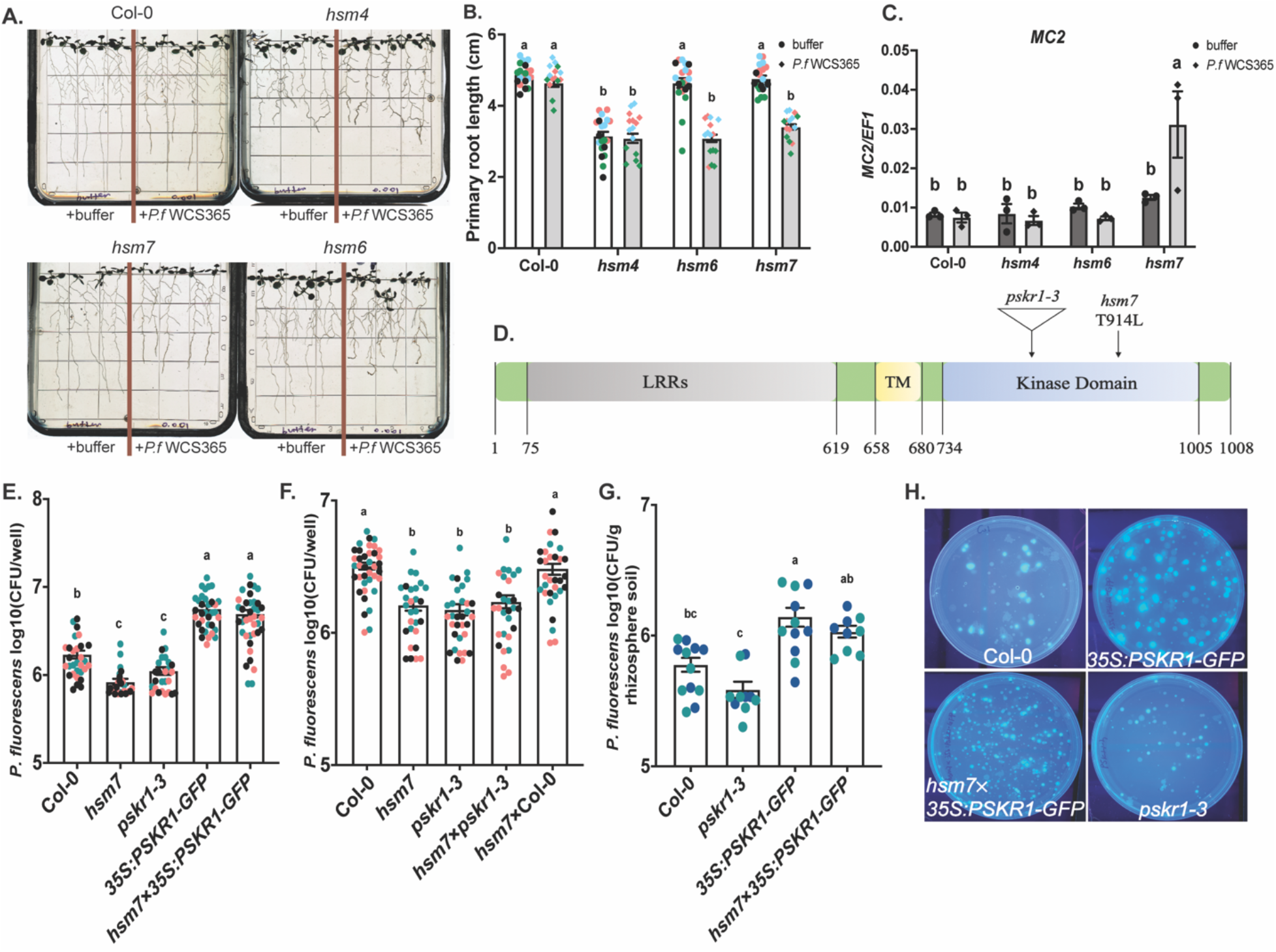
PSKR1 prevents autoimmunity against *P. fluorescens* in the rhizosphere microbiome. **(A)** Root phenotypes of 15-day-old Col-0, *hsm4, hsm6* and *hsm7* seedlings treated with 10 mM MgSO_4_ buffer or *P. fluorescens* WCS365. **(B)** Primary root length of plants shown in (A). **(C)** Root expression levels of *MC2* in Col-0, *hsm4, hsm6* and *hsm7* as measured by qRT-PCR. Expression values were normalized to the expression of house-keeping gene *EF1*. n = 3 replicates with 10 plants per replicate. **(D)** A diagram showing PSKR1 protein domains, the T-DNA insertion location in *pskr1-3* and the point mutation location in *hsm7* confirmed by Sanger sequencing. The point mutation in *hsm7* leads to a predicted amino acid change from Threonine to Leucine at position 914. **(E)** Complementation test result showing *P. fluorescens* colonization of Col-0, *hsm7, pskr1-3, 35S:PSKR1-GFP* and the complemented line *hsm7* × *35S:PSKR1-GFP*. **(F)** An allelism test showing *P. fluorescens* colonization levels of Col-0, *hsm7, pskr1-3*, F1 crosses of *hsm7* × *pskr1-3* and *hsm7* × Col-0. **(G)** Quantification of *P. fluorescens* CFU per gram of rhizosphere soil from wildtype Col-0, *pskr1-3, 35S:PSKR1-GFP* and the complementation line *hsm7* × *35S:PSKR1-GFP* growing in natural soil. Rhizosphere soil from two plants was collected together as one sample. Each dot indicates result from one plant in (B), (E), and (F); each dot represents one sample pooled from two plants in (G). Letters denote significant differences by ANOVA and Tukey’s HSD (p < 0.05) in (B), (C), (E), (F), and (G). Bars indicate means ± s.e.m. Data shown are from three independent experiments colored by experimental replicate. **(H)** Images of King’s B media plates plated with Col-0, *35S:PSKR1-GFP, hsm7×35S:PSKR1-GFP* and *pskr1-3* rhizosphere soil samples normalized to sample weight.

To identify the causative mutation in *hsm7*, we used bulk segregant analysis based on the *hsm* phenotype and next-generation sequencing (Methods). We successfully identified an increase in SNPs frequency on chromosome 2 linked to the *hsm7* phenotype, among which we identified a T914L amino acid substitution in the intercellular kinase domain of the plant gene PHYTOSULFOKINE RECEPTOR 1 (PSKR1) (Figure 1D). As *hsm7* showed enhanced resistance to the root fungal pathogen *Fusarium oxysporum*^9^, and previous studies have found that a loss of *PSKR1* results in enhanced resistance to *F. oxysporum^10^*, we hypothesized that this mutation in *PSKR1* might underlie the *hsm7* phenotype.

To determine if the T914L amino acid substitution in *PSKR1* is the causative mutation, we tested the rhizosphere *P. fluorescens* WCS365 colonization level of a T-DNA insertion mutant *pskr1-3* and a *PSKR1* overexpression line *35S:PSKR1-GFP*. While *pskr1-3* phenocopies *hsm7* with lower rhizosphere *P. fluorescens* levels, *35S:PSKR1-GFP* harbours two-fold higher *P. fluorescens* colonization than wildtype plants (Figure 1E), revealing that *PSKR1* indeed acts as a novel positive regulator that enriches rhizosphere *P. fluorescens* abundance. We found that overexpression of *PSKR1* in the *hsm7* mutant complements the *hsm7* phenotype (Figure 1E) and that the F1 generation of a *hsm7* × *pskr1-3*, but not a *hsm7* × Col-0, cross maintains *hsm7-like P. fluorescens* levels (Figure 1F), indicating that a loss-of-function of *PSKR1* underlies the *hsm7* deficiency in rhizosphere *P. fluorescens* growth.

PSKR1 is a plant receptor-like kinase that perceives the endogenous plant peptide phytosulfokine (PSK)^11,12^, which has been described as a growth-related peptide that promotes root and shoot growth through enhanced cell elongation^13^. Consistent with these previous observations, we found that *pskr1-3* and *hsm7* have shorter primary roots whereas *35S:PSKR1-GFP* shows significantly longer roots than wild-type plants (Extended Figure 1A, 1B). To test if the decreased rhizosphere *P. fluorescens* levels of *pskr1-3* and *hsm7* are due to their smaller root architecture, we quantified *P. fluorescens* rhizosphere colonization in natural soil system (Extended Figure 1C, Methods) which allows quantifying *P. fluorescens* colonies per gram of rhizosphere by normalizing *P. fluorescens* levels to root weight. We found that *35S:PSKR1-GFP* had significantly enriched *P. fluorescens* relative to Col-0 and the *pskr1-3* mutant (Figure 1G, 1H). A *hsm7×35S:PSKR1-GFP* cross also showed significantly higher levels of *P. fluorescens* relative to the *pskr1-3* mutant. This result confirms that *PSKR1* regulates plant association with commensal bacteria and enriches rhizosphere *P. fluorescens* abundance in a natural environment, independent of changes in root architecture.

### PSKR1 specifically enriches *Pseudomonas* abundance in the rhizosphere microbiome

To determine if the effects of PSKR1 on the microbiome are limited to *P. fluorescens* or if they result in community-wide changes, we conducted 16S rRNA sequencing for wildtype plants and *pskr1* loss-of-function mutants *hsm7* and *pskr1-3*. In this experiment, we grew plants in a natural soil mix for 20 days (Methods), harvested and sequenced the plant rhizosphere microbiome. We did not observe significant differences in microbiome community abundance or diversity between Col-0 and *PSKR1* loss-of-function mutants, as measured by alpha-diversity indices including community richness (Species Observed and Chao1) and diversity (Simpson and Shannon) (Figure 2A, Extended Table 1). Similarly, unconstrained principal coordinate analysis (PCoA) showed largely overlapping communities from Col-0, *pskr1-3* and *hsm7* rhizosphere soil samples indicating that *PSKR1* does not affect overall microbiome structure (Figure 2B, Extended Table 2). ANCOM analysis (Figure 2C, Extended Table 3) showed that although most bacteria taxa remain at the same richness level across genotypes, only the Pseudomonadaceae family showed a significant decrease in *pskr1-3* compared with Col-0. By displaying the relative abundance of all ASVs belonging to the Pseudomonadaceae family (Figure 2D, Extended Table 4), we observed that *hsm7* and *pskr1-3* showed reduction of all *Pseudomonas* ASVs relative to Col-0. These results suggest that PSKR1 shapes the rhizosphere microbiome structure by specifically enriching Pseudomonadaceae in the rhizosphere microbiome community without leading to microbiome dysbiosis.

**Figure 2.**
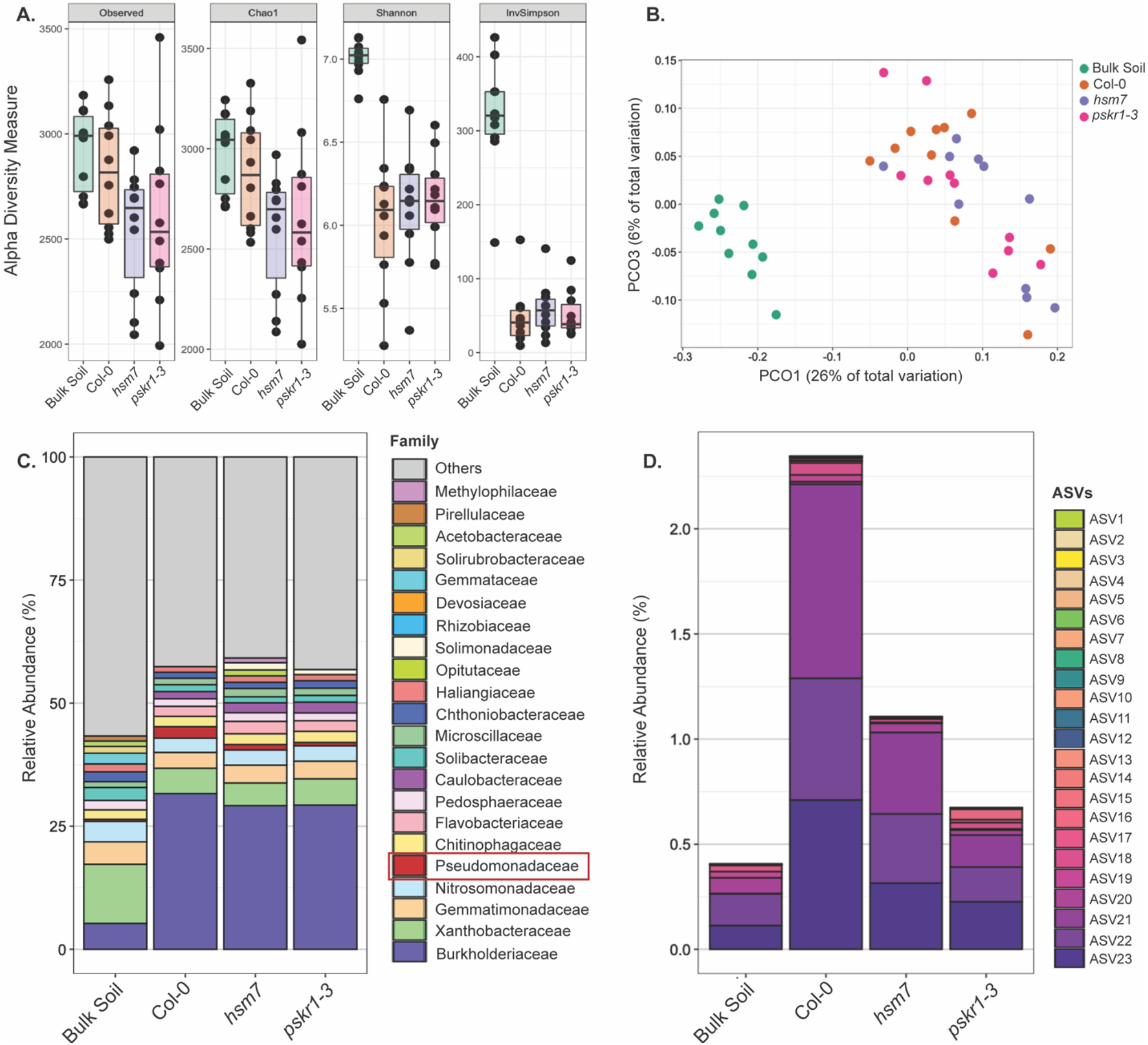
PSKR1 specifically enriches Pseudomonadaceae in the rhizosphere microbiome. (**A**) Chao1, Shannon and Simpson indexes obtained from 16S rRNA sequencing of DNA from bulk soil and Col-0, *hsm7* and *pskr1-3* rhizospheres grown in natural soil for 17 days. In the box plots, the centre line represents the median; the box edges show the first and third quartiles; n=10. (**B**) Principal coordinate analysis (PCoA) based on the Bray-Curtis index of bacterial amplicon sequence variants; n=10. (**C**) Relative abundance of bacteria at the family level in bulk soil and rhizosphere soil samples. (**D**) Relative abundance of Pseudomonadaceae ASVs in bulk soil and rhizosphere soil samples.

### *PSKR1* maintains the plant growth-defense trade-off in response to commensal *P. fluorescens*

We observed that *pskr1* mutants displayed autoimmune-like stunted root growth in the presence of the growth-promoting commensal *P. fluorescens* (Figure 1A, 1B, Extended Figure 2A, 2B), suggesting that *pskr1* mutants may have an inappropriately high defense responses to rhizosphere microbiota. To investigate the mechanisms by which PSKR1 regulates the plant growth-defense trade-off to microbiota, we conducted RNA-sequencing (RNAseq) and analyzed the PSKR1-mediated transcriptome changes in roots. In these experiments, we inoculated Col-0 and *pskr1-3* seedling roots with or without *P. fluorescens* WCS365 and isolated root RNA 2 days after bacterial treatment (Methods).

In wildtype plants, we identified 289 differentially expressed genes (DEGs) upon *P. fluorescens* treatment [adjusted *p* value (*P*adj < 0.05), fold change (FC) ratio (|log_2_FC| ≥1)] including 280 upregulated genes and 9 downregulated genes (Extended Table 5), among which 54 DEGs (18.7% of the total DEGs) were assigned to the GO term defense response [GO:0006952] (Extended Table 5). In the *pskr1-3* mutant, we identified 280 DEGs responding to *P. fluorescens* [162 out of 280 DEGs (57.9%) overlapped with DEGs in Col-0, Extended Table 6], including 270 induced genes and 10 suppressed genes, among which 104 DEGs (37.1% of the total DEGs) belonging to GO term defense response [GO:0006952]. These data indicate that the overall magnitude of *P. fluorescens-* mediated transcriptional changes was similar among plant genotypes, but that *P. fluorescens* induces a distinct transcriptional signature in *pskr1-3* that is more defense-focused than wildtype Col-0 roots.

Next, in order to classify the transcriptional signatures induced by commensal bacteria and determine how they are altered in the *psrk1-3* mutant, we built a protein-protein interaction network (PPI) of the *P. fluorescens*-induced DEGs with STRING. For wildtype plants, we observed a well-integrated bacterial-induced transcriptome network which comprises two subnetworks identified by k-means clustering (Figure 3A, Extended Table 7). Genes belonging to the most significant Gene Ontology (GO) term “cell wall organization (GO:0071555)” and “defense response (GO:0006952)” formed the two subclusters, which are mainly composed of root hair-specific proteins and plant immunity regulators respectively (Extended Table 7). This observation nicely aligns with the known root hair and lateral root growth promotion effects of *P. fluorescens* WCS365^14^ and importantly suggests that plant immune responses are induced to limit commensal bacteria overgrowth in the rhizosphere. For PPI the network analysis of *pskr1-3* mutant, we observed a robust shift of the *pskr1-3* transcriptome network compared with wildtype, where the *pskr1-3* mutant has a larger and denser defense cluster but smaller growth cluster (Figure 3B, Extended Table 8). This observation suggests a shift of focus from growth to defense in the *pskr1-3* mutant and thus a stronger defense response induced by *P. fluorescens* compared with wildtype plants.

**Figure 3.**
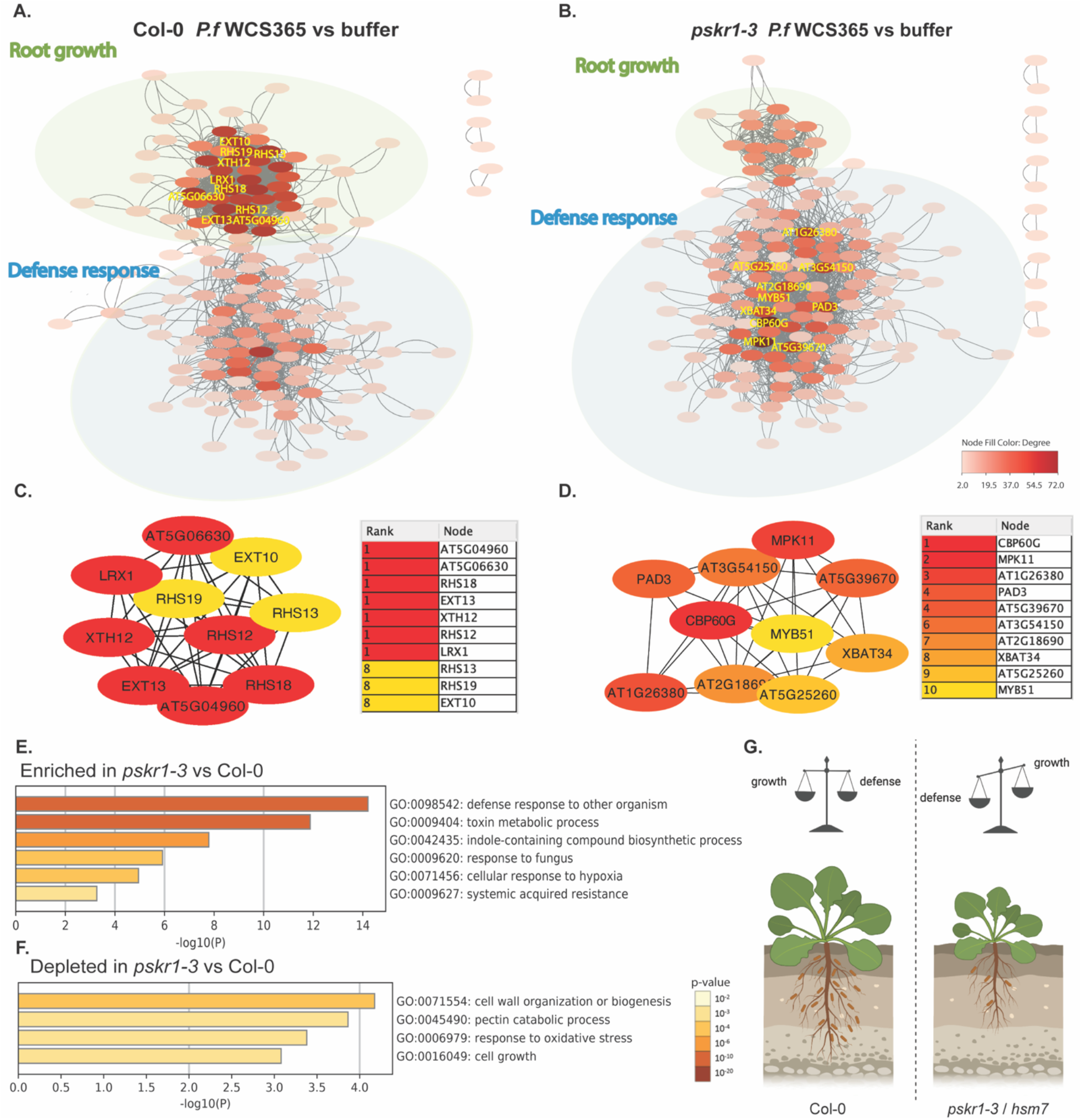
PSKR1 balances the plant growth-defense trade-off in response to beneficial microbes. **(A)(B)** Protein-protein interaction (PPI) network of upregulated DEGs by *P. fluorescens* WCS365 inoculation compared to mock treatment in Col-0 and *pskr1-3* plants. Each node represents a protein, and interactions with a confidence score > 0.4 (above the medium confidence level) are shown. Node fill color represents the node degree. Nodes form two functional clusters according to known protein interconnections, labelled as root growth and defense response respectively according to GO term analysis. Nodes that are recognized as the top 10 hubs (Methods) are labelled with specific gene names. **(C)(D)** Network and list of the top 10 hubs in (B) and (C). Hubs are recognized and ranked according to the node degree. Node filling color indicates the ranking of the individual hub. **(E)(F)** GO term analysis of *P. fluorescens* WCS365 induced DEGs that are positively **(E)** or negatively **(F)** enriched in *pskr1-3* versus Col-0. Bar length and fill color represent *p* values for individual GO term. **(G)** In wildtype plants, PSKR1 enables the plant to maintain a growth-defense balance and helps plant recruit *Pseudomonas* to the rhizosphere microbiome. However, in *pskr1* mutant plants, loss-of-function of *PSKR1* results in autoimmunity in response to microbiota, with stunting, defense gene induction and reductions in rhizosphere *Pseudomonas*.

To further compare the transcriptional signatures induced by *P. fluorescens* in Col-0 and *pskr1-3* mutant plants, we identified PPI network hubs (nodes with a high number of PPI connections to other DEGs) as potential key regulators of *P. fluorescens*-induced plant transcriptome changes. The top 10 mostly highly interconnected hubs derived from *P. fluorescens* induced DEGs in wildtype Col-0 plants contain root hair development-related proteins (RHS12, RHS13 and RHS19), root cell wall organization-related proteins (EXT10, EXT13), and a cell wall biosynthesis protein (XTH12) (Figure 3C), supporting root growth promotion centered transcriptional changes mediated by *P. fluorescens*. However, the top 10 hubs in *pskr1-3* plants are composed of known plant defense response activators, including CBP60G, MYB51, MPK11 and PAD3, revealing a defense-centered autoimmune-like transcriptional profile in the *pskr1-3* mutant (Figure 3D). Similarly, the Gene Ontology (GO) analysis further suggested that compared with Col-0, *pskr1-3* is enriched with *P. fluorescens*-induced DEGs involved in defense responses but lacks those related to cell wall organization and cell growth (Figure 3E, 3F). Collectively, these data indicate that *PSKR1* is necessary to facilitate wildtype plant growth promotion and inhibit inappropriate defense in the presence of growth-promoting rhizosphere bacteria (Figure 3G).

To investigate the specific biological processes regulated by PSKR1 during commensal colonization, we performed the gene set enrichment analysis (GESA) with PSKR1-regulated GO terms by comparing *P. fluorescens* treated *35S:PSKR1-GFP* versus *pskr1-3* datasets *(35S:PSKR1-GFP*_WCS365 versus *pskr1-3*_WCS365). In this analysis, plotted GO terms are clustered according to their similarity. We found that PSRK1-regulated GO terms formed four functional groups (defense, growth, photosynthesis, secondary metabolism; Extended Figure 3A). Consistent with the network analysis results (Figure 3), PSKR1 upregulates growth-related GO term clusters such as “photosynthesis” and “sugar transport”, but downregulates defense-related clusters, including “defense response”, “response to hydrogen peroxide” and “glucosinolate biosynthesis” [DEGs from two representative GO terms defense response (GO:0006952) and photosynthesis (GO:0015979) are displayed in heatmaps; Extended Figure 3B, 3C]. Furthermore, we found that PTI (PAMP-triggered immunity) components (RLKs, BAK1/BKK1), MAPK, calcium signaling pathway and ETI (effector-triggered immunity) components are inhibited by PSKR1 (Extended Figure 4A). As visualized by a ridgeline plot (Extended Figure 4B), PSKR1 may also be involved in regulating several plant metabolism KEGG processes contributing to growth or defense output. Collectively these analyses reveal a critical role for PSKR1 in balancing the plant growth-defense trade-off by regulating multiple growth and defense processes.

### PSKR1 suppresses plant SA-mediated defense gene expression to regulate immunity against rhizosphere microbiota

Consistent with expression of canonical SA responsive genes in our RNAseq dataset *(CBP60G* and *PAD3*), the *pskr1-3* mutant exhibited elevated SA-mediated defenses responses in leaves^15^. Thus, we hypothesized that PSKR1 may inhibit plant SA-mediated defense signaling to avoid autoimmunity against the microbiome. To test the hypothesis, we compared transcriptional profiles of Col-0, *pskr1-3* and *35S:PSKR1-GFP* to SA treated roots (Figure 4A). We found that genes induced by root SA treatment (Figure 4; Cluster B) were strongly inhibited in *35S:PSKR1-GFP*, were induced in the *pskr1-3* mutant, and further induced by *P. fluorescens* treatment. In contrast, genes repressed by SA (Figure 4; Cluster D) were strongly induced in *35S:PSKR1-GFP*, showed lower expression in the *pskr1-3* mutant, and were further repressed by *P. fluorescens* treatment. We validated expression of the SA-responsive gene *SARD1* by RT-qPCR in an independent experiment (Figure 4B) and found that the induction of *SARD1* by *P. fluorescens* treatment is higher in *pskr1-3* and *hsm7* but reduced in *35S:PSKR1-GFP* compared with Col-0. Collectively these data suggest that PSKR1 suppresses *P. fluorescens-induced* SA transcriptional changes in roots.

**Figure 4.**
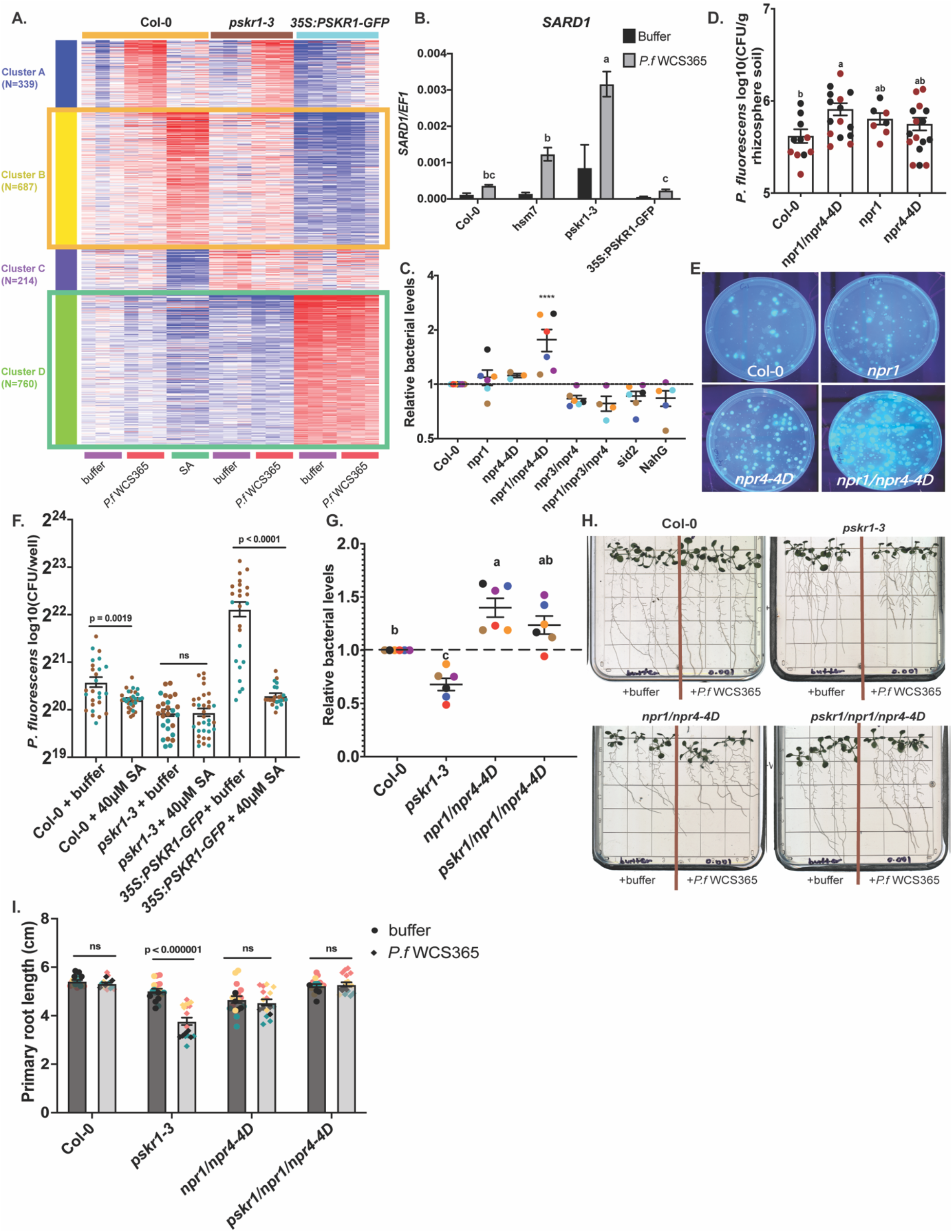
PSKR1 suppresses SA-mediated defenses to enrich rhizosphere *P. fluorescens*. **(A)** Heatmap displaying the top 2000 variable genes from the RNAseq dataset of Col-0, *pskr1-3*, *35S:PSKR1-GFP* plants treated with MgSO4 buffer or *P. fluorescens* for 48 hours, and Col-0 plants treated with 40 μM SA for 6 hours. Cluster B indicates SA-upregulated genes and Cluster D indicates SA-downregulated genes. **(B)** Expression levels of *SARD1* in Col-0, *hsm7, pskr1-3 and 35S:PSKR1-GFP* as measured by qRT-PCR. Expression values were normalized to the expression of house-keeping gene *EF1*. n = 3 replicates with 10 plants per replicate. **(C)** Relative rhizosphere *P. fluorescens* WCS365 levels of the indicated genotypes normalized to Col-0. Each dot represents the mean result from 20-24 plants. **(D)** Quantification of *P. fluorescens* colonies per gram of natural rhizosphere soil samples of Col-0, *npr1/npr4-4D, npr1* and *npr4-4D*. Data from three independent experiments are shown. Bars indicate means ± s.e.m. **(E)** Images of King’s B plates from samples shown in (D). **(F)** The rhizosphere *P. fluorescens* WCS365 levels of Col-0, *pskr1-3* and *35S:PSKR1-GFP* plants with or without 40 μM SA pre-treatment. One dot represents the result from one plant. **(G)** The relative rhizosphere *P. fluorescens* WCS365 levels of the indicated genotypes normalized to Col-0. Each dot represents the mean result from 20-24 plants. ANOVA with Tukey’s HSD were used to determine significance (p < 0.05) in (B), (C), (D), and (G). Different letters (a, b, c and d) or ***(*p* < 0.0005) are used to label genotypes with statistical differences. Bars indicate means ± s.e.m. **(H)** Root phenotype of 15-day-old Col-0, *pskr1-3*, *npr1/npr4-D* and *pskr1/npr1/npr4-4D* seedlings with MgSO4 buffer or *P. fluorescens* WCS365 treatment. **(I)** Primary root length of 15-day-old Col-0,*pskr1-3, npr1/npr4-D* and *pskr1/npr1/npr4-4D* seedlings with MgSO4 buffer or *P. fluorescens* WCS365 treatment. A Student’s t-test was used to test for significance (p<0.05) in (F) and (I). Bars indicate means ± s.e.m. Data from each temporal replate is labelled with different colors in (C), (D), (F), (G) and (I).

To investigate whether plant SA-mediated defenses shape the rhizosphere microbiome, we tested SA mutants for their rhizosphere *P. fluorescens* levels (Figure 4C). These mutants included SA perception mutants (*npr1, npr3/npr4, npr1/npr3/npr4, npr4-4D*, and *npr1/npr4-4D*) and SA deficient genotypes (NahG and *sid2)*. Although most genotypes did not show significantly altered rhizosphere *P. fluorescens* colonization compared with wildtype Col-0, the *npr1/npr4-4D* mutant, which is completely deficient in SA perception^16^, showed significantly higher levels of rhizosphere *P. fluorescens*, suggesting that SA signaling inhibits rhizosphere commensal colonization. Similarly, in natural soil, we observed that the *npr1/npr4-4D* mutant had significantly enriched *P. fluorescens* abundance in the rhizosphere (Figure 4D, 4E). Consistent with the well-studied SA signaling mechanisms in leaves^16^, *npr1* and *npr4-4D* had an additive effect in regulating *P. fluorescence* (Figure 4D). Collectively, our results suggest that SA signaling inhibits *P. fluorescens* abundance in the *Arabidopsis* rhizosphere in the presence of a complex microbiome.

### The PSKR1-mediated autoimmunity against *P. fluorescens* is dependent on SA signaling

To assess if microbiota-triggered elevated defense response in *pskr1* mutants is SA-signaling dependent, we tested whether exogenous SA treatment could affect *P. fluorescens* colonization in the rhizosphere of Col-0 and *pskr1* mutants (Figure 4F). In these experiments, we grew plants hydroponically in 48-well plates with liquid MS media and pre-treated plants with or without 40 μM salicylic acid for 24 hours to induce SA-mediate defense responses before *P. fluorescens* WCS365 inoculation. SA pre-treatment suppressed commensal bacteria colonization in wildtype Col-0, consistent with our results that SA-mediated defense responses inhibit *P. fluorescens* colonization in the rhizosphere (Figure 4C, 4D and 4E). Interestingly, *pskr1-3* was SA-insensitive and did not show significantly reduced *P. fluorescens* colonization after SA pre-treatment. By contrast, SA pre-treatment of *35S:PSKR1-GFP* reduced *P. fluorescens* colonization to wildtype levels, suggesting that the increased *P. fluorescens* colonization in the *35S:PSKR1-GFP* rhizosphere might be due to SA deficiency. Collectively these data indicate that SA modulates the rhizosphere microbiome in a PSKR1-dependent manner.

To test if *pskr1*-mediated autoimmunity against microbiota are wholly or in part through SA signaling, we generated a triple *pskr1/npr1/npr4-4D* mutant and compared the transcriptome to *pskr1-3* and *npr1/4D* mutants. Thorough RNAseq, we found that the increased expression level of SA-inducible genes (Col-0_SA vs. Col-0_buffer) in *pskr1-3* (Cluster A; Extended Figure 5A) was inhibited in both *npr1/npr4-4D* and *pskr1-3/npr1/npr4-4D* mutants. The result was further validated by RT-qPCR with the SA-induced gene *WRKY51* and *WRKY70^16^* (Extended Figure 6), where relative to wildtype plants, the *npr1* and *npr4-4D* mutants had partial decreases in *P. fluorescens-* triggered *WRKY51* and *WRKY70* gene induction, and the gene induction was fully blocked in the *npr1/npr4-4D* and *pskr1-3/npr1/npr4-4D* mutants. These results suggest that the upregulation of SA-dependent gene expression in *pskr1-3* is fully NPR1/NPR4 dependent, and that *pskr1/npr1/npr4-4D* harbours a *npr1/npr4-4D-like* defense gene expression profile.

We then tested if the reduction in bacterial growth and plant stunting in *pskr1* mutants are due to elevated SA signaling in response to *P. fluorescens* inoculation. As is shown in Figure 4G, a *pskr1/npr1/npr4-4D* mutant rescued the rhizosphere *P. fluorescens* WCS365 colonization deficiency of *pskr1-3* to *npr1/npr4D*-like levels demonstrating that PSKR1 functions upstream of SA perception to regulate rhizosphere microbiota. Consistently, while *pskr1-3* shows significant root stunting in the presence of *P. fluorescens* WCS365 (Figure 4H, 4I), neither *npr1/npr4D* nor *pskr1/npr1/npr4-4D* displayed a stunted autoimmune phenotype with commensal bacteria treatment, indicating that *pskr1-3* autoimmunity in response to commensal bacteria is fully SA dependent. These data indicate that PSKR1 suppresses plant SA-mediated defense responses to maintain a plant growth-defense balance during associations with the rhizosphere microbiome.

Our results thus far demonstrate that PSKR1 balances plant growth-defense trade-off during host-microbiome association, and that the higher defense output in the *pskr1* mutants is due to elevated SA-mediated defenses. To determine if all *PSKR1-* dependent downstream signaling events, such as growth and metabolism processes (Extended Figure 2, Extended Figure 3), are SA-dependent, we also investigated other transcriptional changes regulated by PSKR1. As is shown in Extended Figure 5A, genes that are not SA-inducible (Col-0_SA vs. Col-0_buffer) but are induced (Cluster B) or suppressed (Cluster C) by the *pskr1* mutation (*pskr1-3*_buffer vs. Col-0_buffer) show intermediate expression in the *pskr1-3/npr1/npr4-4D* triple mutant between the expression levels in *pskr1-3* and *npr1/npr4-4D*, suggesting that the expression of those PSKR1-regulated genes are not solely NPR3/NPR4 dependent. GO term analysis (Extended Figure 5B) for Cluster B and C suggests that PSKR1 regulates multiple plant growth and metabolism-related pathways that are not fully SA-dependent. These clusters include positively PSKR1 regulated GO terms related to iron homeostasis (Cluster C), which are associated with maintaining photosynthesis and chloroplast integrity^17^, and negatively PSKR1 regulated GO terms related to response to hypoxia stress (Cluster B), which is known to trigger ROS homeostasis and acclimation^18^. Collectively, these data suggests that PSKR1 may regulate multiple processes in addition to SA-mediated defense responses in the presence of rhizosphere microbiota to balance the plant growth-defense trade-off.

### *P. fluorescens* upregulates PSKR1 expression to manipulate host responses

The results that *PSKR1* recruits *P. fluorescens* to the plant rhizosphere raises the question of whether commensal bacteria could upregulate plant *PSKR1* expression to hijack plant signaling and promote their own rhizosphere colonization. Using an *Arabidopsis PSKR1pro:GUS* reporter line, we observed a strong induction of *GUS* expression in roots treated with either *P. fluorescens* WCS365 bacteria or bacterial culture supernatant (Figure 5). We found that heat-killed *P. fluorescens* bacteria could no longer induce *PSKR1* expression. However, heat-killed bacterial supernatant still strongly upregulated *PSKR1* expression level in the root, which suggests that *P. fluorescens* may secrete a small molecule or peptide that induces *PSKR1* expression. This indicates that *P. fluorescens* may be able to manipulate *PSKR1* activity to promote its own recruitment to the rhizosphere.

**Figure 5.**
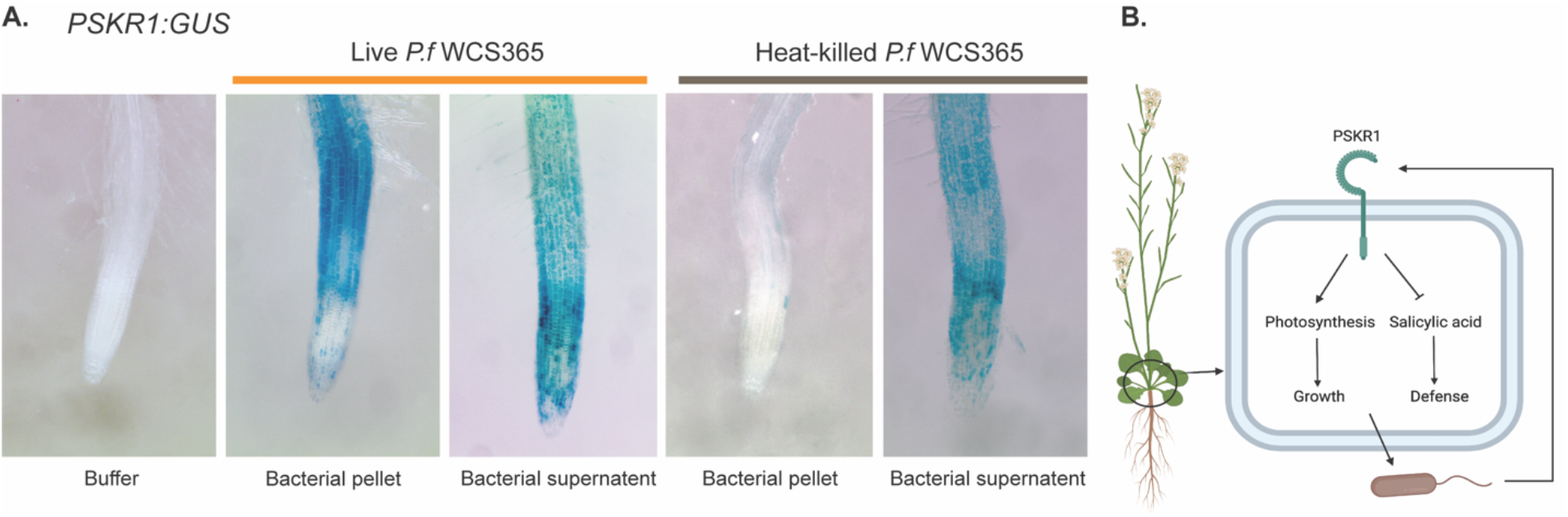
*P. fluorescens* WCS365 upregulates *PSKR1* expression in roots. (A) *PSKR1pro:GUS* plants were inoculated with buffer, *P. fluorescens* WCS365 (OD_600_ = 0.002) or culture supernatant, or heat-treated *P. fluorescens* (OD_600_ = 0.2) or culture supernatant for 24 hours. (B) A working model for the role of PSKR1 in recruiting *P. fluorescens. P. fluorescens* upregulates root *PSKR1* expression to hijack plant signaling. The activation of *PSKR1* in turn recruits more *P. fluorescens* by upregulating photosynthesis and other growth-related events while suppressing SA-mediated defense. Through fine-tuning plant growth-defense trade-off, PSKR1 helps plant recruit *Pseudomonas* to the rhizosphere.

## Discussion

Beneficial microbiota can promote plant growth and improve plant fitness; however, in the presence of growth-promoting microbiota, plants must also mount a sufficient defense response to avoid parasitism by the microbiome. This work demonstrates a novel genetic mechanism underlying the largely uncharacterized fundamental question of how plants avoid autoimmunity and balance growth-defense trade-off in response to the microbiota. We show that through suppressing SA-mediated root defenses and upregulating multiple growth-related processes, PSKR1 fine-tunes plant energy distribution between defense and growth, shapes the rhizosphere microbiome, and enables beneficial *Pseudomonas* to better colonize the plant rhizosphere. We found that *P. fluorescens* could, in turn, upregulate root *PSKR1* gene expression to potentially manipulate plant signaling and promote its own colonization in the rhizosphere.

While previous work has extensively characterized plant autoimmune mutants that are hyper-immune-sensitive to pathogen attack at the expense of growth^4^, how plants avoid autoimmunity to rhizosphere microbiota is largely unexplored. Compared with many well-characterized autoimmune mutants with elevated SA levels that are constructively stunted^4^, the autoimmunity phenotype of *pskr1* mutants is largely specific to treatment with commensals. This highlights the importance of maintaining the appropriate level of plant defense while coordinating commensal-induced growth: too much defense can result in plant autoimmunity to commensals; too little may result in bacteria overgrowth and susceptibility to pathogens. Consistently, we found the *35S:PSKR1:GFP* overexpression line had increased levels of rhizosphere commensals, and has previously been shown to have enhanced susceptibility to biotrophic foliar pathogens^15^. Collectively, our findings demonstrate the importance of the PSKR1 signaling pathway in balancing plant defense levels to rhizosphere microbiota with growth costs, helping plants to coordinate with microbiota-mediated growth promotion while surviving under the microbe-enriched natural environment.

Plant sulfated peptides are widely conserved in the plant kingdom^19^, suggesting that using PSKR1 signaling to balance the growth-defense trade-off in association with rhizosphere microbiota might be a common mechanism across different plant taxa. Recent work by Ogawa-Ohnishi *et al*.^20^ showed that another closely related plant sulfated peptide family PSY (PLANT PEPTIDE CONTAINING SULFATED TYROSINE) also mediates plant growth-stress tolerance trade-off through cell-to-cell communication. Interestingly, similar to our result showing that *P. fluorescens* induces *PSKR1* expression, the PSY signaling pathway can be manipulated by the rice blast pathogenic bacteria *Xanthomonas oryzae* pv. *oryzae (Xoo)*, which secretes a sulphated RaxX peptide that mimics the PSY1 structure^21^. These results shed light on the importance of plant sulfated peptides as a component of conserved machinery in balancing plant growth and responses to diverse biotic and abiotic stresses. These signaling pathways can be hijacked by pathogens or beneficial microbes to suppress host defense signaling and benefit their proliferation.

Understanding how organisms regulate their immune responses to recruit or suppress microbiota is integral to maintaining host health. As soil *Pseudomonas* abundance is associated with improvements in agricultural production due to its known fungal disease suppression and growth promotion abilities^22,23^, dynamically and specifically enriching *Pseudomonas* could be beneficial for crop health. Our work demonstrates that PSKR1 enriches beneficial *Pseudomonas* abundance in the plant rhizosphere, which suggests that manipulation of plant PSKR1 signaling could be a means to recruit *Pseudomonas*. Engineering of the crop rhizosphere microbiome to enrich beneficial microbes has the potential to enhance plant disease resistance and nutrient uptake in an environmentally-friendly way.

## Methods

### Plant materials and growth conditions

The *hsm* mutants^7^, *pskr1-3^13^, PSKR1:GUS^24^, 35S:PSKR1-GFP^25^*, *npr1/npr4-4D^16^, npr1^26^, npr4-4D^16^, npr3/npr4*^27^, *sid2-2*^28^, NahG^29^ genotypes were previously described. The *pskr1-3/npr1/npr4-4D* triple mutant was generated by crossing *pskr1-3* with *npr1/npr4-4D*. F1 and F2 plants were genotyped through PCR to detect the presence and homozygosity for T-DNA insertions in *PSKR1* as well as SNP mutations in *NPR1* and *NPR4* genes with the primers listed in Extended Table 9.

*Arabidopsis* seeds were surface sterilized by chlorine gas for 1 hour and stored at 4°C for 2 days before planting. Plants were grown in growth rooms or growth chambers at 23°C under 12h light/12h dark or 16h light/8h dark with light intensity at approximately 80 μmol m^-2^ s^-1^. For the 48-well plate colonization assay and GUS staining assay, plants were grown hydroponically in 48-well plates with 300 μl Murashige and Skoog (MS) with 2% (w/v) sucrose media liquid in each well. For natural soil colonization assays, plants were grown in natural soil mix (described below) with the natural soil collected from UBC farm (49°15.0’N, 123°14.4’W), Vancouver, British Columbia, Canada. For RT-qPCR experiments, root growth phenotype quantification and RNAseq experiments, *Arabidopsis* plants were grown vertically on ½X MS with 2% (w/v) sucrose solid media plates for 5 days and transferred to ½X MS solid media for two days before inoculation.

### Bacteria inoculation and root phenotype quantification

To observe root growth after inoculation with commensal bacteria *P. fluorescens* strain WCS365, we first grew *Arabidopsis* plants vertically on half-strength MS with 2% (w/v) sucrose media for five days and transplanted healthy plants to half-strength MS media for two days. For bacterial inoculations, the overnight culture of *P. fluorescens* WCS365 was resuspended with 10 mM MgSO4 buffer and diluted to a final OD600 0.001. 10 μL of the inoculum or 10 mM MgSO4 buffer was pipetted along the root of 7-day-old *Arabidopsis* seedlings. After 7 days of growth after inoculation, the plants were then scanned, and the primary root length was then measured using ImageJ^30^.

### Commensal bacteria colonization quantification assay

We used two different methods to quantify commensal bacteria abundance in the plant rhizosphere: the 48-well plate hydroponic system and the natural soil system as previously described^8,31^. Briefly, for the 48-well plate assay, sterilized *Arabidopsis* seeds were plated on Teflon mesh disks (Mcmaster Carr) floating on 300 μL full strength MS media with 2% sucrose in each well of the 48-well plate. After 10-days growing hydroponically under 16h light/8h dark conditions, the media was replaced with 270 μL fresh half-strength MS media for two days, and the 12-day plant roots were inoculated with the luciferase-tagged *P. fluorescens* strain WCS365-Luc^8^ at a final OD_600_ 0.00002. The luminescence signal was measured by the plate reader two days after inoculation and plotted onto a standard curve to estimate bacterial CFU.

For the natural soil colonization system, natural soil containing natural microbiota was collected from the UBC farm (49°15.0’N, 123°14.4’W), Vancouver, British Columbia, Canada as previously described^8^. The collected soil was sieved with 3 mm-pore sieves,homogenized and mixed with calcine clay (Turface) and perlite [1:0.5:1 natural soil: calcine clay (Turface): perlite] to improve drainage and plant growth. 6-day-old *Arabidopsis* seedlings grown on ½X MS plates with 2% sucrose were then transplanted to the natural soil mix. After 17 days of growing at 12h light/12h dark with light intensity 80 μmol m^-2^ s^-1^ in a growth chamber, the rhizosphere soil samples were harvested as described^8^. In brief, we gently shook the plant to loosen the soil particles attached to the root until just the most closely adhered soil remained. Rhizosphere samples from two plants of the same genotype were collected in one tube as one biological replicate. To normalize samples to soil and root weight, we weighed samples and added buffer (7.5 mM MgSO4 and 20% glycerol) to a final concentration of 0.05 g soil/mL. Samples were homogenized with a Tissuelyser (2 minutes at 30 rps) and serially diluted with buffer (7.5 mM MgSO_4_ and 20% glycerol) to a final concentration of 0.00025 g/mL. 100 μL of the diluted rhizosphere soil samples was plated on King’s B media, after which *P. fluorescens* colonies were identified under a UV light source and counted.

### Bulk segregation and mapping of *hsm7*

The *HSM7* (PSKR1) gene was mapped as previously described^32^. Briefly, *hsm7* was outcrossed to Ler to generate F1 hybrid, which was self-segregated to produce a segregating F2 population. A total of 65 F2 plants (30 homozygous and 35 heterozygous) were selected based on root GUS reporter activity following COR and fig22 co-treatment. Genomic DNA from the 65 selected F2 plants were isolated, pooled, subjected to library preparation and paired-end sequencing with Illumina Hi-seq. With the whole-genome sequencing analysis protocol as described^32^, an enrichment of Col-0 SNPs, that is corresponding to the enrichment of the *hsm7* mutation, was observed on chromosome 2 spanning a region of 2.9 million base pairs to delimit the causal *hsm7* mutation. Further sequence analyses revealed a single candidate gene with a ‘G’ to ‘A’ nucleotide change in the coding region of the *PSKR1* gene, which was further confirmed by Sanger sequencing.

### RNA isolation and gene expression

For RNAseq and qRT-PCR experiments, *Arabidopsis* plants were grown vertically on ½X MS with 2% (w/v) sucrose solid media plates for 5 days, after which 10 healthy-looking seedlings were transplanted to one fresh half-strength MS media plate for 2 days before inoculation. For bacterial inoculations, overnight cultures of *P. fluorescens* strain WCS365 were resuspended in 10 mM MgSO4 buffer and diluted to OD_600_ 0.01. 10 μL of bacteria inoculum, 10mM MgSO4 or 40 μM SA was pipetted along the root of 7-day-old *Arabidopsis* seedlings on half-strength MS solid media. Root samples of 10 plants from the same media plate were bulk harvested as one biological replicate. Samples were collected 48 hours after bacterial treatment and 6 hours after SA treatment, snap-frozen in liquid nitrogen and stored at −80°C until RNA extraction.

For qRT-PCR experiments, RNA was extracted with the RNeasy Plant Mini Kit (Qiagen) and quantified by Nanodrop. 800 ng of RNA for each sample was reverse-transcribed to cDNA through Oligo(dT)-primed reverse transcription with Invitrogen™ SuperScript™ III Reverse Transcriptase (Fisher Scientific). The qPCR was performed with Applied Biosystems™ PowerUp™ SYBR™ Green Master Mix (Fisher Scientific) with the housekeeping gene *EF1* as an internal control. The primers were previously described^7,16^ and listed in Extended Table 9.

### RNA sequencing and analysis

RNA was isolated with the RNeasy Plant Mini Kit (Qiagen) and submitted to Michael Smith Genome Sciences Center (http://www.bcgsc.ca/) for library construction and sequencing. Paired-end 75 bp RNA-sequencing was performed using an Illumina Hi-seq 2500 platform. The adapter sequence was trimmed with bbduk (version 38.86)^33^ and filtered with trimmomatic (version 0.39)^34^. High-quality reads were mapped to the Tair10 genome from Tair^35^ (https://www.arabidopsis.org/) and quantified with STAR (version 2.7)^36^. Differential expression analysis was performed in R with the DESeq2^37^ package. Network analysis (Figure 3A, 3B) was done with STRING^38^; the network hubs (Figure 3D, 2E) were identified with cytoHubba^39^ in Cytoscape^40^. Significant GO terms for the gene set enrichment analysis (GESA) (Extended Figure 3A) were identified with g:Profiler, clustered with EnrichmentMap^41^ and annotated with AutoAnnotate^42^ in Cytoscape. The Heatmaps (Figure 4A, Extended Figure 5A) and KEGG pathway diagram (Extended Figure 4A) were generated with iDEP.95^43^. The GO term enrichment analysis (Figure 3E, 3F, Extended Figure 5B) was performed and visualized with Metascape^44^. The heatmaps in GESA analysis (Extended Figure 3B, 3C) and Ridgeline Diagrams (Extended Figure 4B) were generated with NetworkAnalyst^45^.

### GUS staining assay

The GUS staining assay to detect *PSKR1* expression in the root was performed as described^46^. Briefly, *PSKR1:GUS* plants were grown in liquid MS media supplemented with 2% sucrose within 48-well plates (4-6 seedlings per well) for 4 days under 12h light/12h dark condition. The media was then replaced with 270 μL fresh 1/2X MS media for 2 days before inoculation. Inoculum was prepared by centrifuging the *P. fluorescens* WCS365 overnight culture to pellet bacterial cells. The supernatant was filter-sterilized with 0.2 mm-pore syringe filters and the bacterial pellet was resuspended in 10 mM MgSO_4_ to a final OD_600_ 0.02. For the heat-killed bacterial treatment, a *P. fluorescens* WCS365 overnight culture was boiled at 95°C for 15 minutes. After centrifugation, the supernatant was filter-sterilized with 0.2 mm-pore syringe filters; the heat-killed bacterial pellet was resuspended with 10 mM MgSO_4_ to a final OD_600_ 2. For 6-day-old hydroponically grown plants, 30 μL of each inoculum or 10 mM MgSO_4_ was added to the 270 μL 1/2X MS media for a 22-hour treatment in the growth chamber. The media was then replaced with 300 μl GUS staining solution (50 mM pH 7 sodium phosphate buffer, 10 mM EDTA, 0.5 mM potassium ferricyanide, 0.5 mM potassium ferrocyanide, 0.5 mM x-gluc, 0.01% Triton X-100) and incubated for 2 hours at 37°C in the dark before the GUS signal was observed.

### Microbiome 16S rRNA sequencing

Col-0, *hsm7* and *pskr1-3* plants were vertically grown on 1/2X MS with 2% (w/v) sucrose solid media as described above. 6-day-old *Arabidopsis* seedlings were transplanted into a natural soil mix [1:0.5:1 natural soil: calcine clay (Turface): perlite]. After 17 days of growing in natural soil at 12h light/12h dark in the growth chamber, the rhizosphere soil samples and bulk soil samples were harvested as previously described. Rhizosphere samples from four plants of the same genotype were collected in one tube as one biological replicate, which was then frozen in liquid nitrogen and stored at −80°C.

The 16S rRNA sequencing library was prepared as described in the Earth Microbiome Project Illumina 16S rRNA protocol (www.earthmicrobiome.org). Briefly, total DNA was extracted from soil samples with DNeasy PowerSoil Pro Kit (Qiagen). PCR was performed with primers 515F–806R targeting the V4 region of the 16S rRNA using Phusion High-Fidelity DNA Polymerase (NEB). The PCR product concentration was measured with Quant-iT PicoGreen dsDNA Assay Kits (Invitrogen) and normalized for all samples. Paired-end 300 bp sequencing was performed with Illumina MiSeq.

For raw FASTQ files, adapter sequences and primers were removed with cutadapt v.2.1^47^. Trimmed sequences were qualify filtered and processed in QIIME2 version 2019.10^48^. A 515F- and 806R-trained V4 SILVA database 132^49^ was used to classify amplicon sequence variants (ASVs) with Deblur^50^. α- and β-Diversity metrics were calculated by Phyloseq^51^ package in R version 1.4.1717. Using Bray-Curtis dissimilarity, principal coordinate analysis (PCoA) was conducted on Hellinger-transformed data to assess the bacterial community structure distribution (β diversity). Permutational multivariate analysis (PERMANOVA) was performed to identify differences in β diversity distances across groups. Pairwise analysis of microbiome composition (ANCOM) in QIIME2^52^ was applied to determine bacterial taxa abundance among genotypes.

## Data availability

Raw data for 16S rRNA sequencing is available in the Sequence Read Archive database under BioProject PRJNA896256, accession numbers SRR22133746 to SRR22133785. Raw data for RNA sequencing is available in the Sequence Read Archive database under BioProject PRJNA897874, accession numbers SRR22177437 to SRR22177481.

## Acknowledgments

This work was supported by an NSERC Discovery Grant and Accelerator Award (NSERC-RGPIN-2021-03587) and a Canada Research Chair salary award to C.H.H. Additional trainee support was provided by Chinese Scholarship Council Awards to S.S. Early stages of this work were supported by NIH grant R37 GM48707 and NSF grants MCB-0519898 and IOS-0929226 awarded to Frederick M. Ausubel.

---

**Extended Figure 1.**
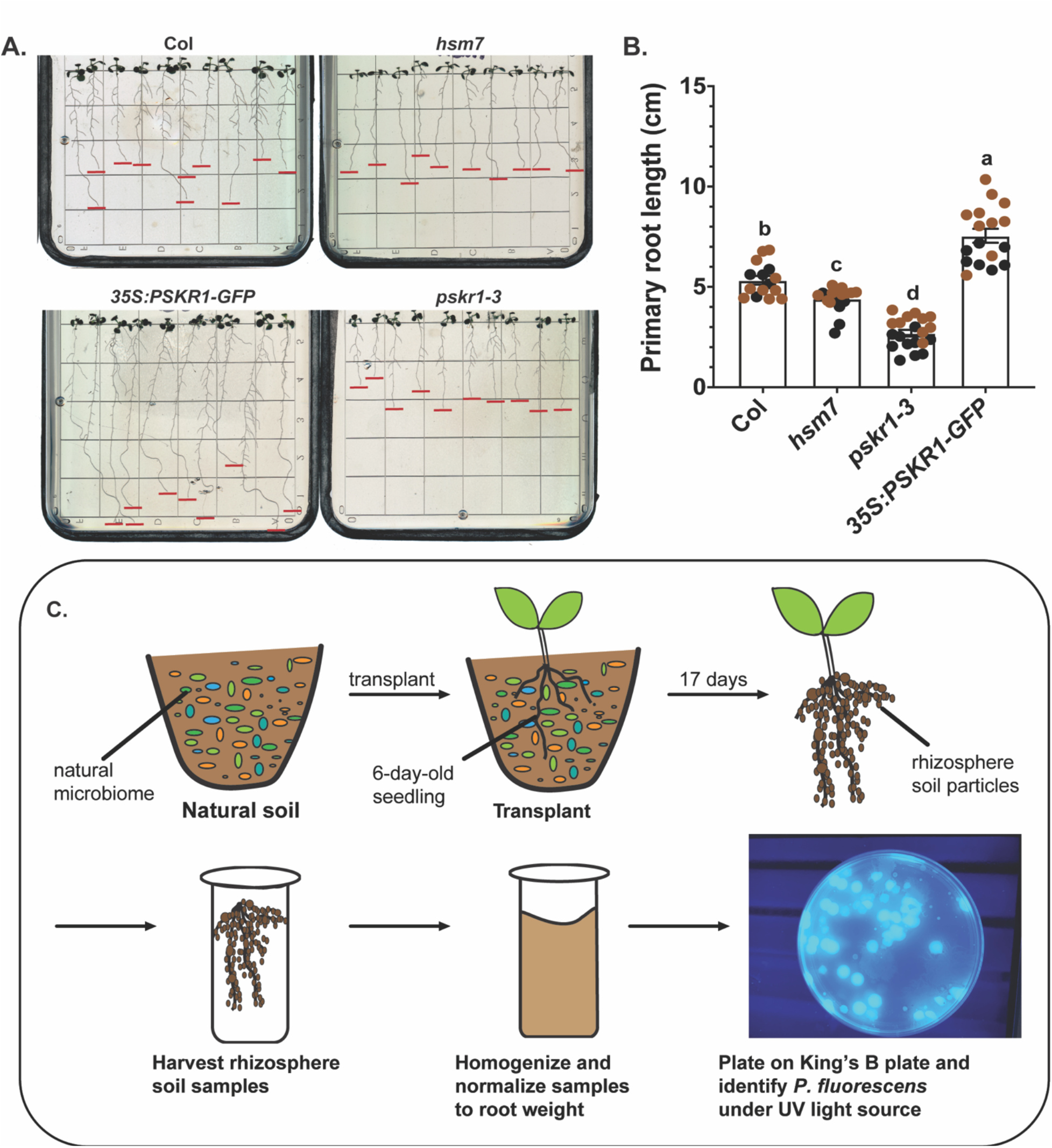
PSKR1 enriches rhizosphere *P. fluorescens* abundance independent of changes in root architecture. (**A**) Root phenotype of 8-day-old Col-0, *hsm7, 35S:PSKR1-GFP* and *pskr1-3* seedlings planted on *1/2* MS media plate with 2% sucrose. The red lines highlight the ending point of the primary roots. (**B**) Primary root length of 8-day-old Col-0, *hsm7,pskr1-3* and *35S:PSKR1-GFP* seedlings. Each dot indicates the primary root length of one plant. Letters denote significant difference by ANOVA and Tukey’s HSD (p < 0.05). Bars indicate means ± s.e.m. Data from two independent experiments are shown and colored by experiment. (**C**) A diagram showing the natural soil colonization system. Natural soil containing natural microbiota was collected from the UBC farm, and 6-day-old plant seedlings from ½ MS plates were transplanted. After 17 days growing under 12h light/12h dark in growth chambers, root rhizosphere soil was collected, weighed and buffer was added to 0.05 g/mL (7.5 mM MgSO4 and 20% glycerol). The homogenized rhizosphere soil in buffer was plated on King’s B media after dilution, after which *P. fluorescens* colonies were identified under an UV light source and counted.

**Extended Figure 2.**
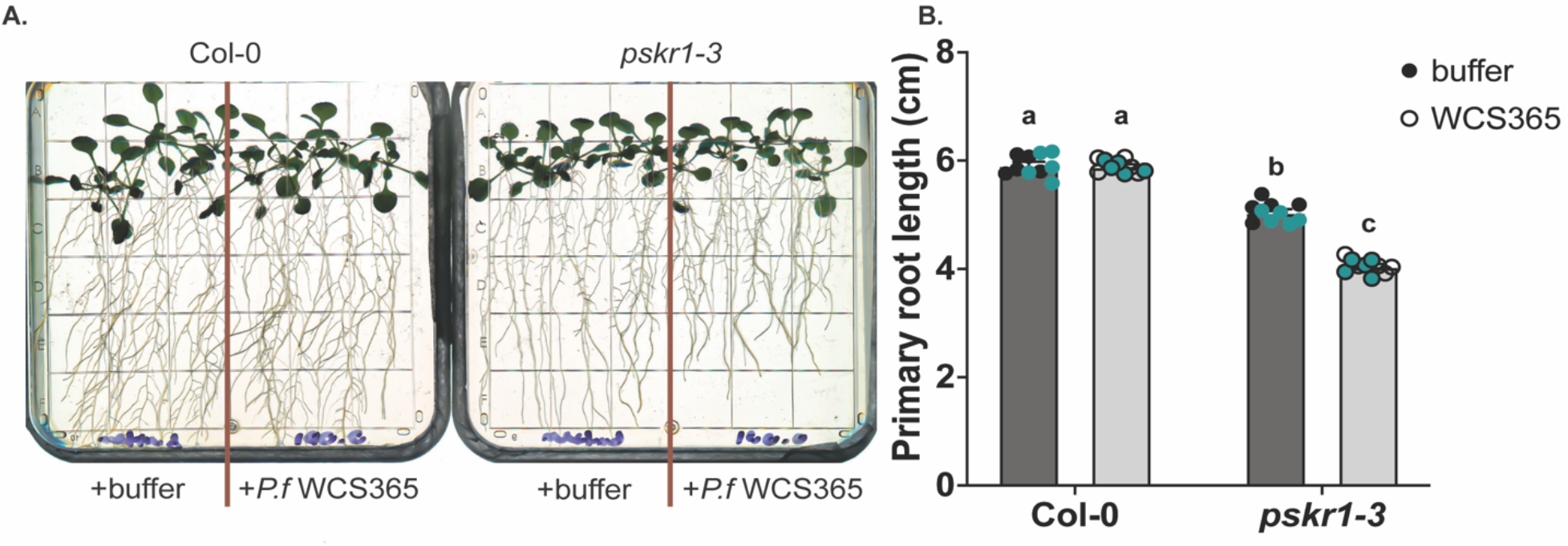
*pskr1-3* shows autoimmune growth deficiency phenotype with commensal *P. fluorescens* treatment. **(A)** Root phenotype of 18-day-old Col-0 and *pskr1-3* seedlings with MgSO4 buffer or *P. fluorescens* WCS365 treatment. **(B)** Primary root length of 18-day-old Col-0 and *pskr1-3* seedlings with MgSO4 buffer or *P. fluorescens* WCS365 treatment. Each dot indicates the primary root length of 1 plant. Letters indicate statistical differences *(p* < 0.05) by ANOVA and Tukey’s HSD. Bars indicate means ± s.e.m. Data from each temporal replate are labelled with different colors.

**Extended Figure 3.**
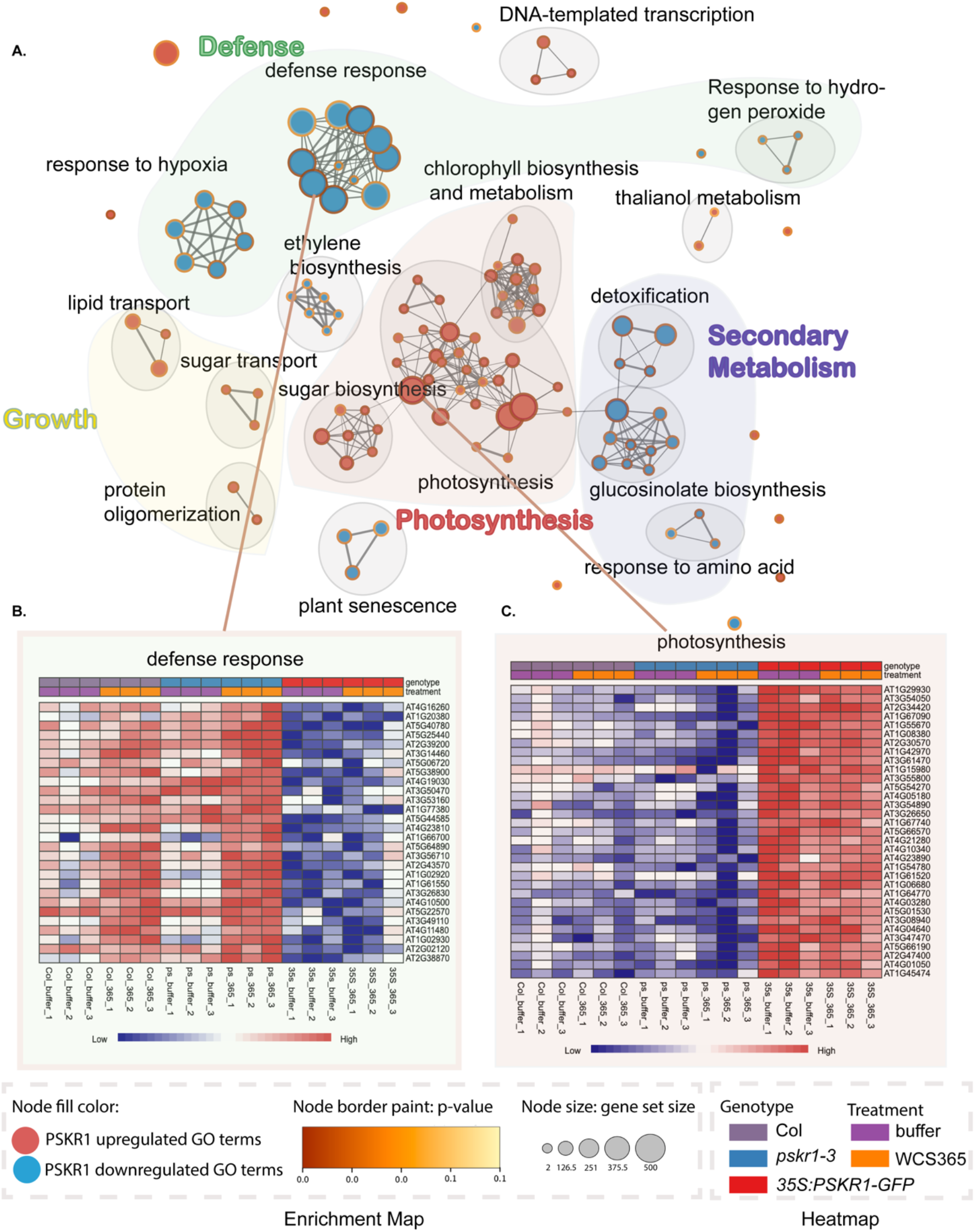
Gene set enrichment analysis (GESA) revealed plant defense or growth-related pathways dependent on PSKR1. **(A)** GESA map of enriched GO terms for DEGs in *35S:PSKR1-GFP* versus *pskr1-3* 48 hours after *P. fluorescens* inoculation *(35S:PSKR1-GFP*_WCS365 versus *pskr1-3_365, P*adj < 0.05, |log_2_FC| ≥ 2). Individual GO terms are plotted as one node; node size represents the number of genes belonging to each GO term; node border color indicates the GO term significance; node filling color suggests PSKR1 positively (red) or negatively (blue) regulated GO terms. Similar GO terms are clustered and circled; similar clusters form functional groups labelled defense, growth, photosynthesis and secondary metabolism. **(B)(C)** Genes belonging to GO term defense response (GO:0006952) and photosynthesis (GO:0015979) specifically for Col-0, *pskr1-3, 35S:PSKR1-GFP* samples with or without *P. fluorescens* treatment are displayed in heatmaps.

**Extended Figure 4.**
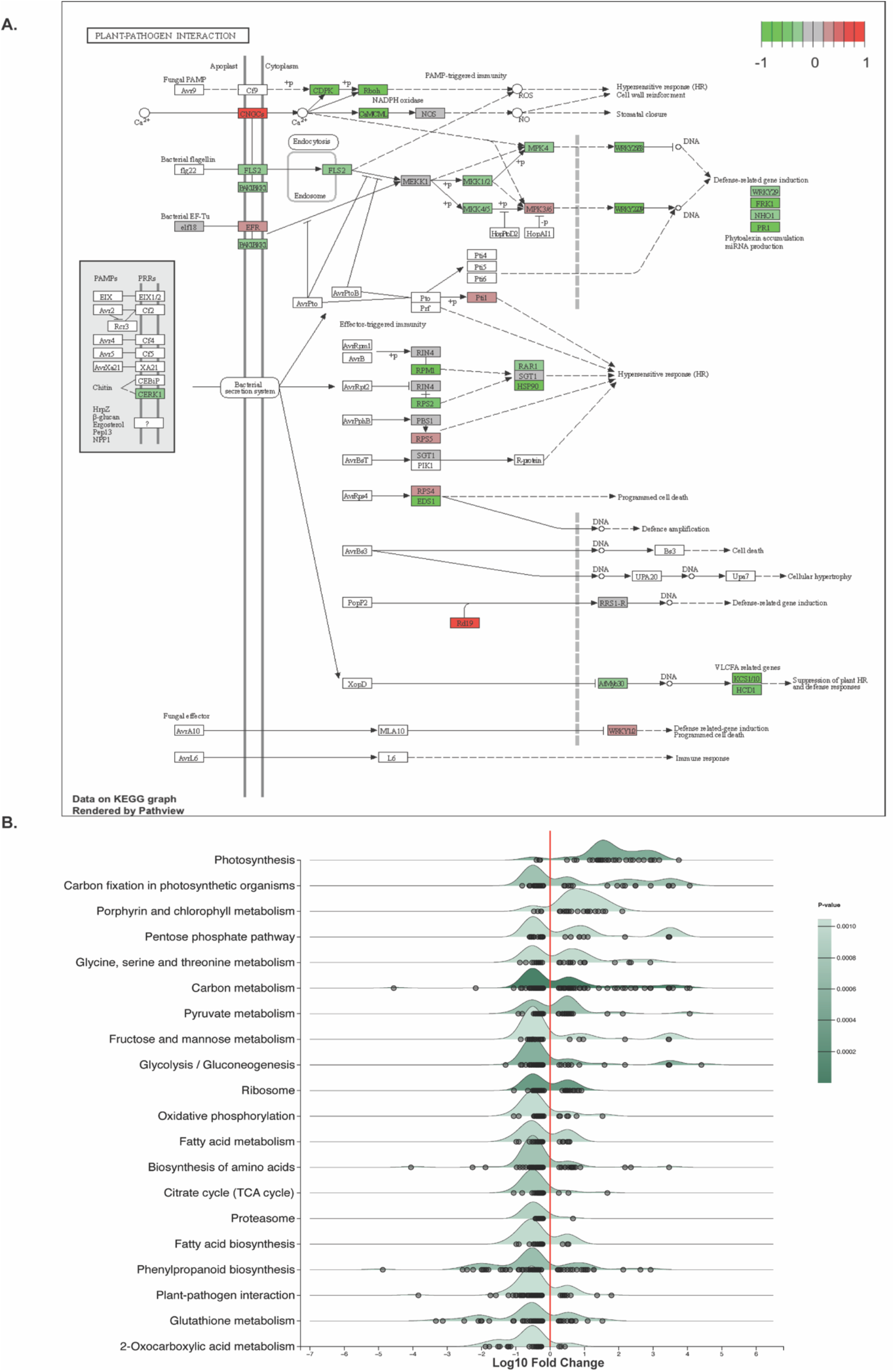
PSKR1 upregulates growth and suppresses defense responses. (**A**) KEGG search and color pathway analysis of PSKR1-regulated DEGs *(35S:PSKR1-GFP_WCS365* versus *pskr1-3*_WCS365, adjusted *p* value (*P*adj < 0.05))in the plant-pathogen interaction pathway by comparing *P. fluorescens* WCS365 treated *35S:PSKR1-GFP* versus *pskr1-3* samples. Red and green color indicate PSKR1 up- and down-regulated DEGs respectively. (**B**) The ridgeline plot visualizing expression distributions of PSKR1 regulated DEGs with the presence of commensals *[35S:PSKR1-GFP_WCS365* versus *pskr1-3_WCS365*, adjusted *p* value (*P*adj < 0.05)] by comparing *P. fluorescens* WCS365 treated *35S:PSKR1-GFP* versus *pskr1-3* samples. The black circles indicate the Log_10_ fold change of individual DEG in each KEGG pathway.

**Extended Figure 5.**
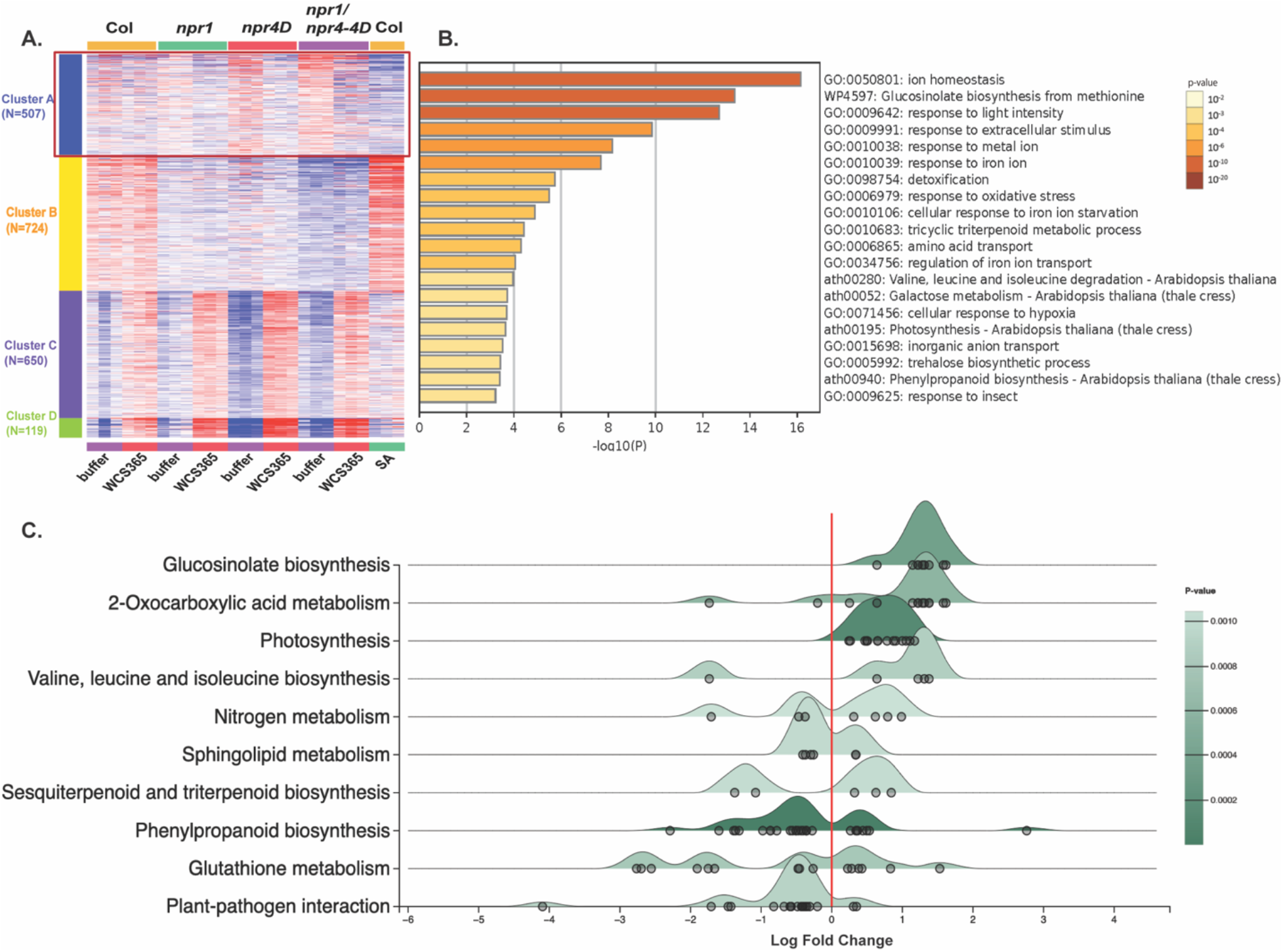
NPR3/4 are involved in regulating plant metabolic processes. (**A**) Heatmap displaying the top 2000 variable genes from the RNAseq dataset of Col-0, *npr1, npr4-4D*, and *npr1/npr4-4D* plants treated with MgSO4 buffer or *P. fluorescens* for 48 hours, and Col-0 plants treated with 40 μM SA for 6 hours. Cluster A indicates SA-downregulated genes that are alleviated by *npr4-4D* mutation. (**B**) The Gene Ontology (GO) processes enriched in Cluster A genes by Metascape. (**C**) The ridgeline plot visualizing expression distributions of NPR3/4-regulated DEGs [*npr4-4D*_buffer versus Col-0_buffer, adjusted *p* value (*P*adj < 0.05)] by comparing buffer treated *npr4-4D* versus Col-0 samples. The black circles indicate the Log_10_ fold change of individual DEGs in each KEGG pathway.

**Extended Figure 6.**
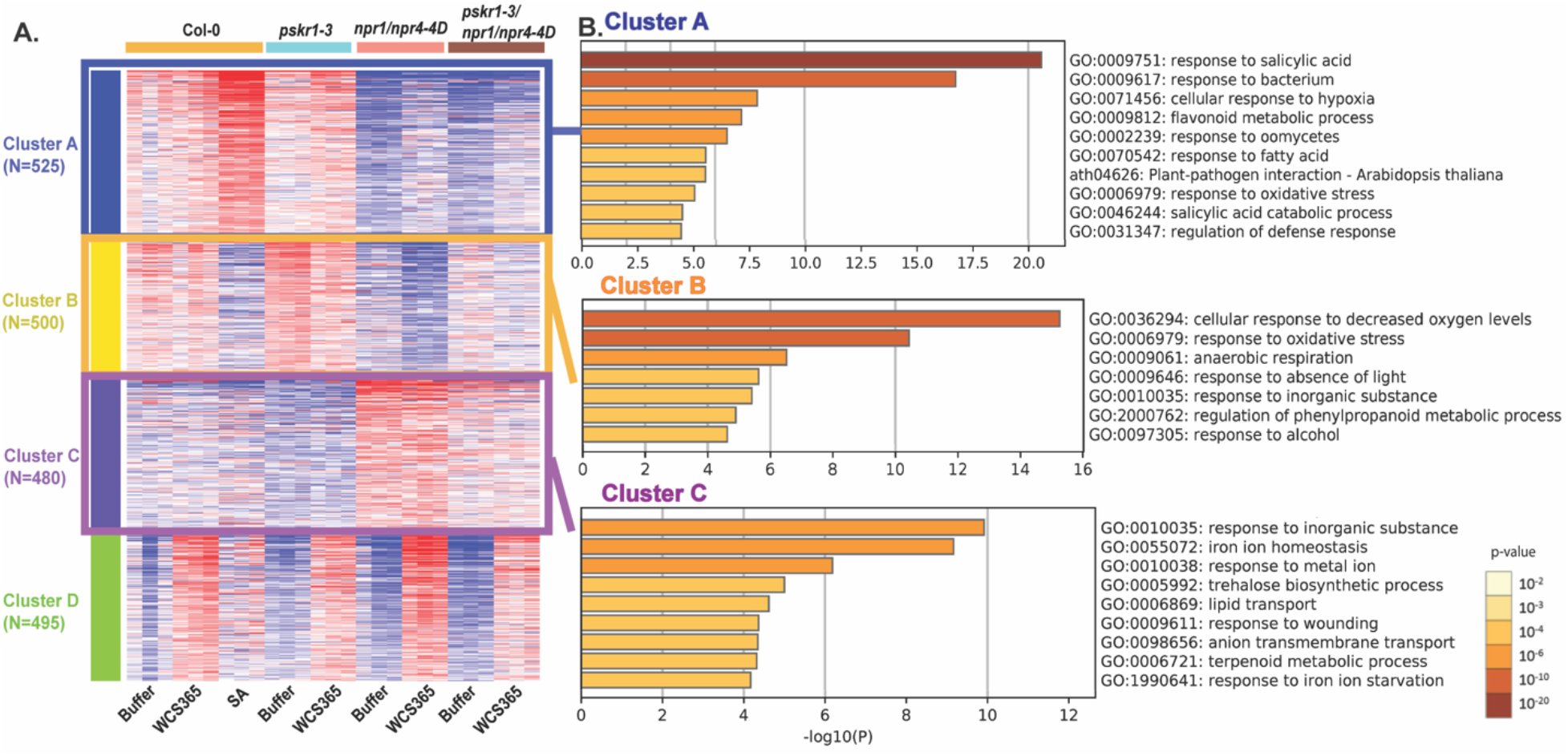
PSKR1 regulates multiple processes in addition to SA-mediated defense responses. (**A**) Heatmap displaying the top 2000 variable genes from the RNAseq dataset of Col-0, *pskr1-3, npr1/npr4-4D*, and *pskr1-3/npr1/npr4-4D* plants treated with MgSO4 buffer or *P. fluorescens* for 48 hours, and Col-0 plants treated with 40 μM SA for 6 hours. Cluster A indicates SA-upregulated genes (Col-0_SA versus Col-0_buffer) that show epistasis by *npr1/npr4-4D* in the *pskr1-3/npr1/npr4-4D* triple mutant. Cluster B and Cluster C indicate PSKR1-regulated genes without a full epistasis effect by *npr1/npr4-4D*. (**B**) The Gene Ontology (GO) processes enriched in Cluster A, B and C genes by Metascape. (**C**) Expression levels of *WRKY51* in Col-0, *npr1, npr4-4D, npr1/npr4-4D* and *pskr1-3/npr1/npr4-4D* as measured by qRT-PCR. Expression values were normalized to the expression of house-keeping gene *EF1*. n = 3 replicates with 10 plants per replicate.

**Extended Figure 7.**
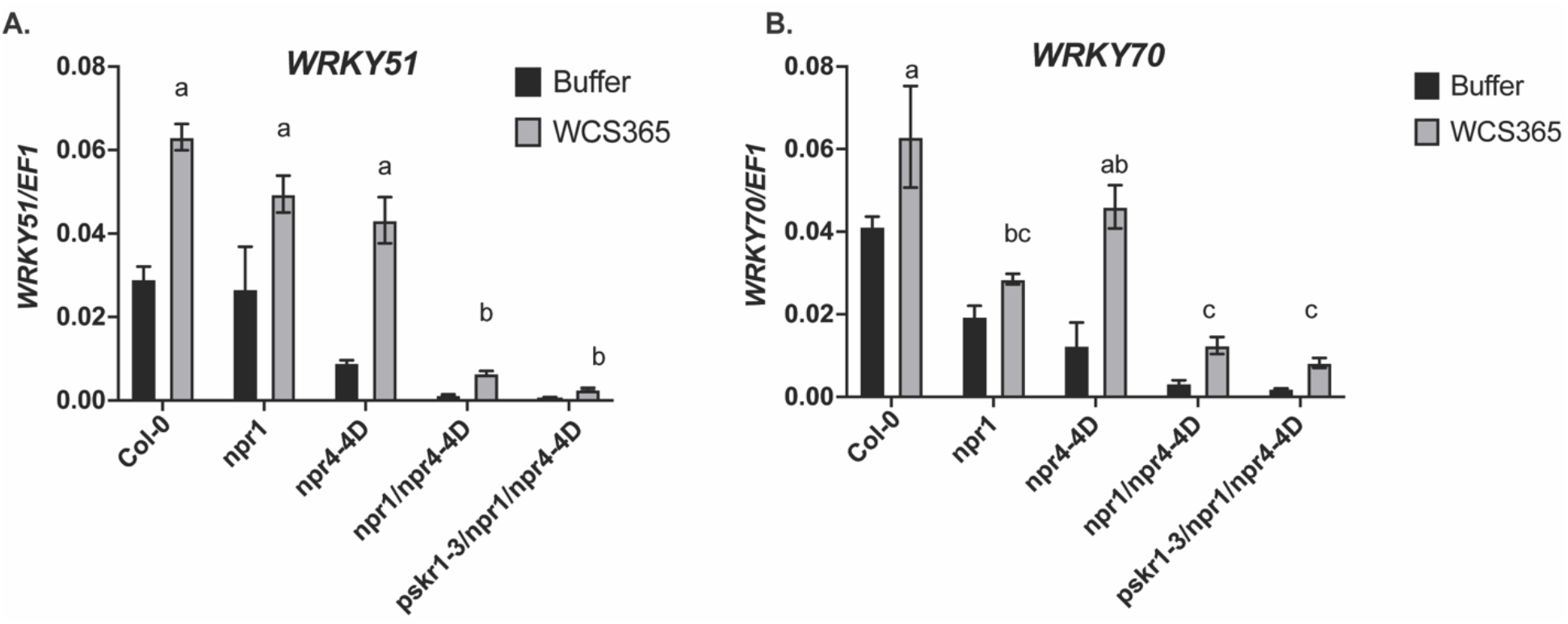
The SA inducible gene expression is inhibited in *pskr1-3/npr1/npr4-4D*. **(A)(B)** Expression levels of *WRKY51* **(A)** and *WRKY70* **(B)** in Col-0, *npr1, npr4-4D, npr1/npr4-4D* and *pskr1-3/npr1/npr4-4D* as measured by qRT-PCR. Expression values were normalized to the expression of house-keeping gene *EF1*. n = 3 replicates with 10 plants per replicate.

## Reference

1. Tringe, S. G. et al. Comparative metagenomics of microbial communities. Science 308, 554–557 (2005).

2. Bakker, P. A. H. M. et al. The Soil-Borne Identity and Microbiome-Assisted Agriculture: Looking Back to the Future. Molecular Plant (2020) doi:10.1016/j.molp.2020.09.017.

3. Chen, T. et al. A plant genetic network for preventing dysbiosis in the phyllosphere. Nature (2020) doi:10.1038/s41586-020-2185-0.

4. van Wersch, R., Li, X. & Zhang, Y. Mighty dwarfs: Arabidopsis autoimmune mutants and their usages in genetic dissection of plant immunity. Front. Plant Sci. 7, 1–8 (2016).

5. Lebeis, S. L. et al. Salicylic acid modulates colonization of the root microbiome by specific bacterial taxa. Science (80-.). 349, 860–864 (2015).

6. Lv, S. et al. Dysfunction of histone demethylase IBM1 in Arabidopsis causes autoimmunity and reshapes the root microbiome. ISME J. (2022) doi:10.1038/s41396-022-01297-6.

7. Zhang, X.-C., Millet, Y. A., Cheng, Z., Bush, J. & Ausubel, F. M. Jasmonate signalling in Arabidopsis involves SGT1b–HSP70–HSP90 chaperone complexes. Nat. Plants 1, (2015).

8. Song, Y. et al. FERONIA restricts Pseudomonas in the rhizosphere microbiome via regulation of reactive oxygen species. Nat. Plants (2021) doi:10.1038/s41477-021-00914-0.

9. Yi Song, Xue-Cheng Zhang, Yichun Qiu, Annika Briggs, Yves Millet, Diana Bartenstein, Catherine Mankiw, Mary L. Cerulli, Jenifer Bush, Keith L. Adams, Andrew C. Diener, Frederick M. Ausubel, C. H. H. A screen for mutants deficient in coronatine-mediated suppression of root immunity identifies. bioRxiv 1–27 (2021) doi: https://doi.org/10.1101/2021.09.12.459990.

10. Shen, Y. & Diener, A. C. Arabidopsis thaliana RESISTANCE TO FUSARIUM OXYSPORUM 2 Implicates Tyrosine-Sulfated Peptide Signaling in Susceptibility and Resistance to Root Infection. PLoS Genet. 9, (2013).

11. Matsubayashi, Y., Shinohara, H. & Ogawa, M. Identification and functional characterization of phytosulfokine receptor using a ligand-based approach. Chem. Rec. (2006) doi:10.1002/tcr.20090.

12. Matsubayashi, Y., Ogawa, M., Kihara, H., Niwa, M. & Sakagami, Y. Disruption and overexpression of Arabidopsis phytosulfokine receptor gene affects cellular longevity and potential for growth. Plant Physiol. (2006) doi:10.1104/pp.106.081109.

13. Hartmann, J., Stührwohldt, N., Dahlke, R. I. & Sauter, M. Phytosulfokine control of growth occurs in the epidermis, is likely to be non-cell autonomous and is dependent on brassinosteroids. Plant J. (2013) doi:10.1111/tpj.12056.

14. Simons, M. et al. Gnotobiotic system for studying rhizosphere colonization by plant growth-promoting Pseudomonas bacteria. Mol. Plant-Microbe Interact. 9, (1996).

15. Mosher, S. et al. The tyrosine-sulfated peptide receptors PSKR1 and PSY1R modify the immunity of Arabidopsis to biotrophic and necrotrophic pathogens in an antagonistic manner. Plant J. 73, 469–482 (2013).

16. Ding, Y. et al. Opposite Roles of Salicylic Acid Receptors NPR1 and NPR3/NPR4 in Transcriptional Regulation of Plant Immunity. Cell (2018) doi:10.1016/j.cell.2018.03.044.

17. Geider, R. J. & La Roche, J. The role of iron in phytoplankton photosynthesis, and the potential for iron-limitation of primary productivity in the sea. Photosynthesis Research vol. 39 (1994).

18. Lai, A. G. et al. Circadian Clock-Associated 1 regulates ROS homeostasis and oxidative stress responses. Proc. Natl. Acad. Sci. U. S. A. 109, (2012).

19. Sauter, M. Phytosulfokine peptide signalling. Journal of Experimental Botany vol. 66 (2015).

20. Peptide, P. & Sulfated, C. Peptide ligand-mediated trade-off between plant growth and stress response. 180, 175–180 (2022).

21. Pruitt, R. N. et al. A microbially derived tyrosine-sulfated peptide mimics a plant peptide hormone. New Phytol. (2017) doi:10.1111/nph.14609.

22. Expósito, R. G., de Bruijn, I., Postma, J. & Raaijmakers, J. M. Current insights into the role of Rhizosphere bacteria in disease suppressive soils. Frontiers in Microbiology (2017) doi:10.3389/fmicb.2017.02529.

23. Lugtenberg, B. & Kamilova, F. Plant-growth-promoting rhizobacteria. Annual Review of Microbiology (2009) doi:10.1146/annurev.micro.62.081307.162918.

24. Stührwohldt, N., Dahlke, R. I., Steffens, B., Johnson, A. & Sauter, M. Phytosulfokine-α controls hypocotyl length and cell expansion in Arabidopsis thaliana through phytosulfokine receptor 1. PLoS One 6, (2011).

25. Hartmann, J., Fischer, C., Dietrich, P. & Sauter, M. Kinase activity and calmodulin binding are essential for growth signaling by the phytosulfokine receptor PSKR1. Plant J. 78, 192–202 (2014).

26. Cao, H., Bowling, S. A., Gordon, A. S. & Dong, X. Characterization of an Arabidopsis Mutant That Is Nonresponsive to Inducers of Systemic Acquired Resistance. Plant Cell 6, 1583–1592 (1994).

27. Zhang, Y. et al. Negative regulation of defense responses in Arabidopsis by two NPR1 paralogs. Plant J. 48, (2006).

28. Wildermuth, M. C., Dewdney, J., Wu, G. & Ausubel, F. M. Isochorismate synthase is required to synthesize salicylic acid for plant defence. Nature 414, 562–565 (2001).

29. Delaney, T. P. et al. A central role of salicylic acid in plant disease resistance. Science (80-.). 266, (1994).

30. Abràmoff, M. D., Magalhães, P. J. & Ram, S. J. Image processing with imageJ. Biophotonics International vol. 11 (2004).

31. Haney, C. H., Samuel, B. S., Bush, J. & Ausubel, F. M. Associations with rhizosphere bacteria can confer an adaptive advantage to plants. Nat. Plants 1, (2015).

32. Zhang, X. C., Millet, Y., Ausubel, F. M. & Borowsky, M. Next-gen sequencing-based mapping and identification of ethyl methanesulfonate-induced mutations in Arabidopsis thaliana. Curr. Protoc. Mol. Biol. 2014, (2014).

33. Bushnell, B. BBMap. https://sourceforge.net/projects/bbmap/ (2015).

34. Bolger, A. M., Lohse, M. & Usadel, B. Trimmomatic: A flexible trimmer for Illumina sequence data. Bioinformatics 30, (2014).

35. Lamesch, P. et al. The Arabidopsis Information Resource (TAIR): Improved gene annotation and new tools. Nucleic Acids Res. 40, (2012).

36. Dobin, A. et al. STAR: Ultrafast universal RNA-seq aligner. Bioinformatics 29, (2013).

37. Love, M. I., Huber, W. & Anders, S. Moderated estimation of fold change and dispersion for RNA-seq data with DESeq2. Genome Biol. 15, (2014).

38. Szklarczyk, D. et al. STRING v11: Protein-protein association networks with increased coverage, supporting functional discovery in genome-wide experimental datasets. Nucleic Acids Res. 47, (2019).

39. Chin, C. H. et al. cytoHubba: Identifying hub objects and sub-networks from complex interactome. BMC Syst. Biol. 8, (2014).

40. Shannon, P. et al. Cytoscape: A software Environment for integrated models of biomolecular interaction networks. Genome Res. 13, (2003).

41. Reimand, J. et al. Pathway enrichment analysis and visualization of omics data using g:Profiler, GSEA, Cytoscape and EnrichmentMap. Nat. Protoc. 14, (2019).

42. Kucera, M., Isserlin, R., Arkhangorodsky, A. & Bader, G. D. AutoAnnotate: A Cytoscape app for summarizing networks with semantic annotations [version 1; referees: 2 approved]. F1000Research 5, (2016).

43. Ge, S. X., Son, E. W. & Yao, R. iDEP: An integrated web application for differential expression and pathway analysis of RNA-Seq data. BMC Bioinformatics 19, (2018).

44. Zhou, Y. et al. Metascape provides a biologist-oriented resource for the analysis of systems-level datasets. Nat. Commun. 10, (2019).

45. Zhou, G. et al. NetworkAnalyst 3.0: A visual analytics platform for comprehensive gene expression profiling and meta-analysis. Nucleic Acids Res. 47, (2019).

46. Millet, Y. A. et al. Innate immune responses activated in Arabidopsis roots by microbe-associated molecular patterns. Plant Cell 22, 973–990 (2010).

47. Martin, M. Cutadapt removes adapter sequences from high-throughput sequencing reads. EMBnet.journal 17, (2011).

48. Bolyen, E. et al. Reproducible, interactive, scalable and extensible microbiome data science using QIIME 2. Nature Biotechnology vol. 37 (2019).

49. Quast, C. et al. The SILVA ribosomal RNA gene database project: Improved data processing and web-based tools. Nucleic Acids Res. 41, (2013).

50. Amir, A. et al. Deblur Rapidly Resolves Single-Nucleotide Community Sequence Patterns. mSystems 2, (2017).

51. McMurdie, P. J. & Holmes, S. Phyloseq: An R Package for Reproducible Interactive Analysis and Graphics of Microbiome Census Data. PLoS One 8, (2013).

52. Mandal, S. et al. Analysis of composition of microbiomes: a novel method for studying microbial composition. Microb. Ecol. Heal. Dis. 26, (2015).

