## Extended Table 1 for "PSKR1 balances the plant growth-defense trade-off in the rhizosphere microbiome"

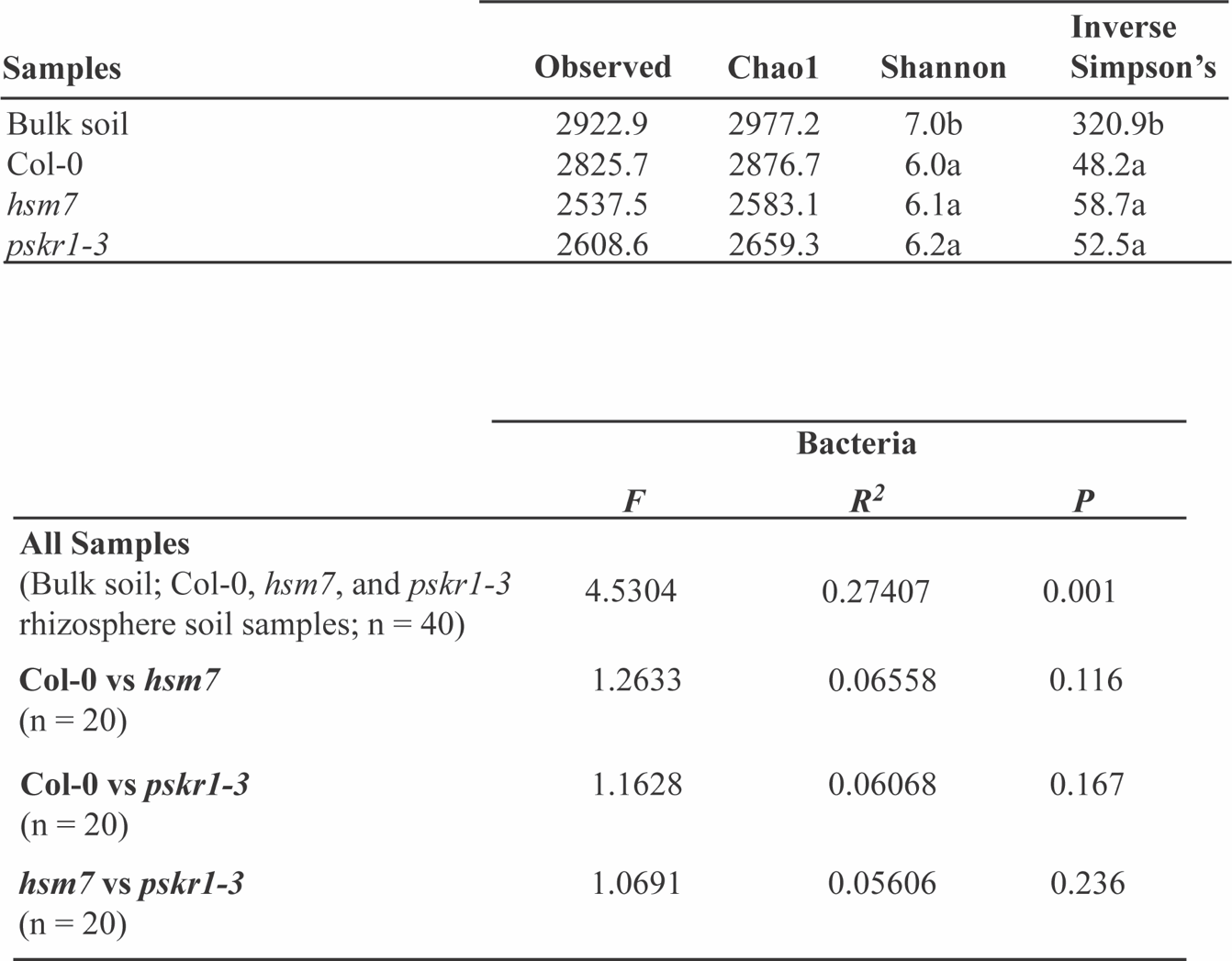

**Extended Table 1. ANOVA and Tukey’s post-hoc test (p < 0.05) of alpha diversity indices for *Arabidopsis* rhizosphere and bulk soil samples.**
