## Extended Table 2 for "PSKR1 balances the plant growth-defense trade-off in the rhizosphere microbiome"

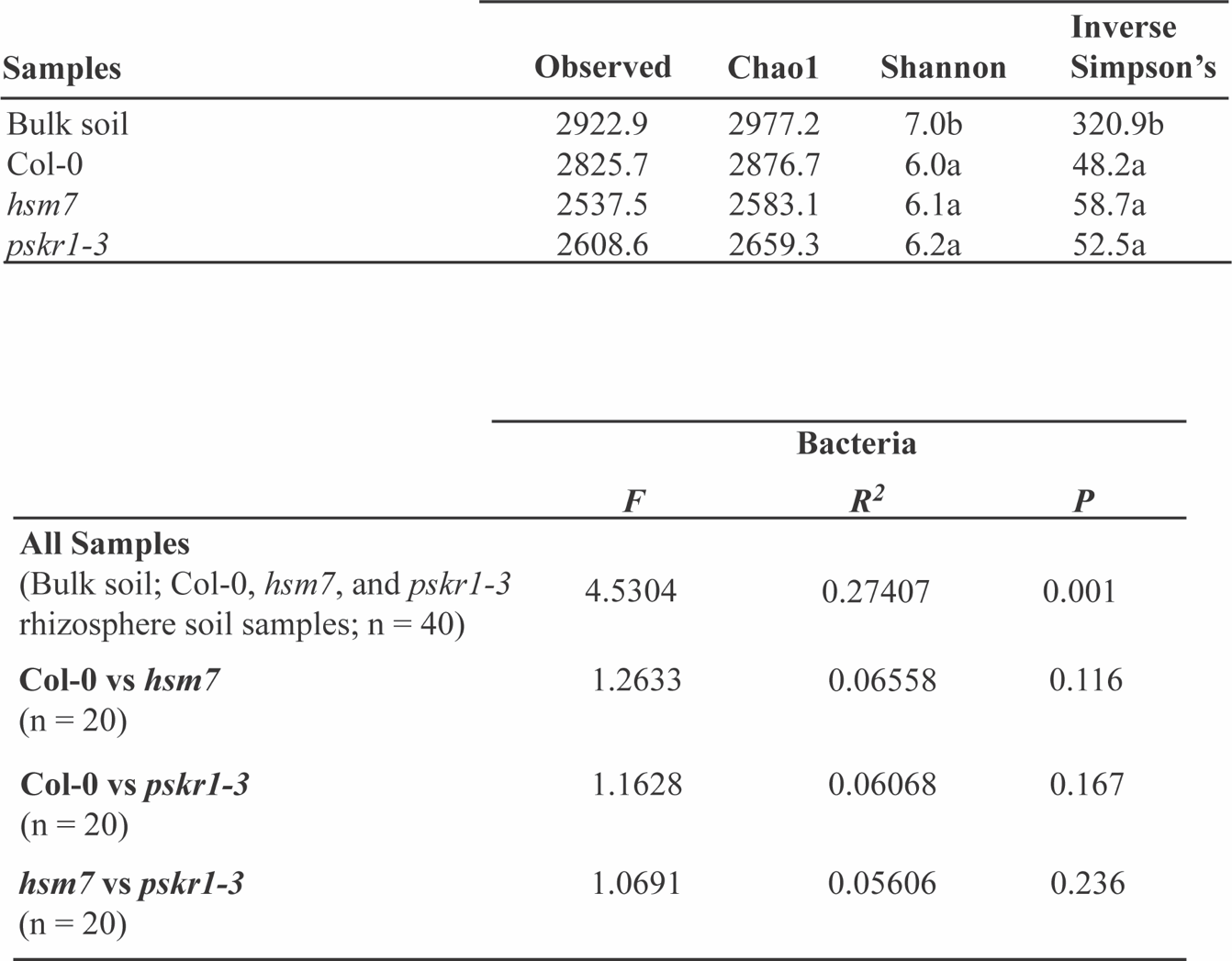

**Extended Table 2. Permutational multivariate analysis of the bacterial communities associated with *Arabidopsis* rhizosphere.**
