## Extended Table 3 for "PSKR1 balances the plant growth-defense trade-off in the rhizosphere microbiome"

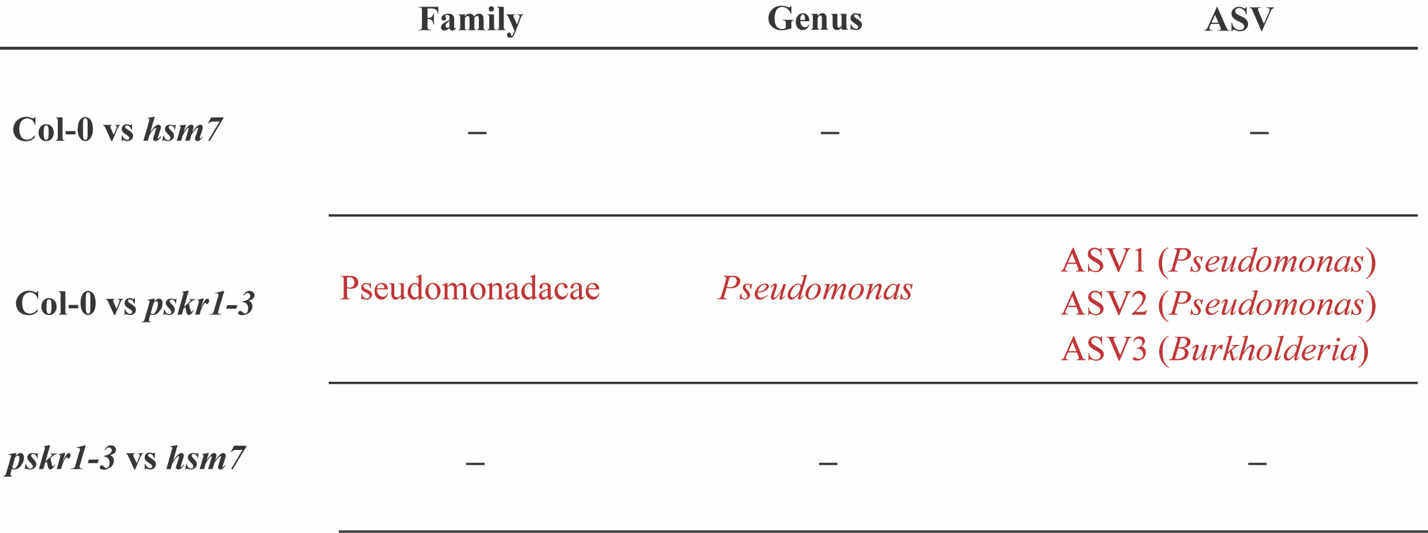

**Extended Table 3. ANCOM test results for significant bacterial differential abundance within *Arabidopsis* genotype comparisons.** Red color indicates significantly enriched bacteria taxa. The 16S rRNA sequences for the enriched ASVs are included in Extended Table 4.
